# Cutting through host autofluorescence: fluorescence lifetime imaging microscopy for visualising intracellular bacteria in Symbiodiniaceae

**DOI:** 10.1101/2024.01.16.575970

**Authors:** Pranali Deore, Sarah Jane Tsang Min Ching, Douglas R. Brumley, Madeleine J.H. van Oppen, Elizabeth Hinde, Linda L. Blackall

## Abstract

- Photoperiodicity is key to the synchronization of life stages in Symbiodiniaceae, *Breviolum minutum* which harbors taxonomically diverse epi- and endosymbiotic bacteria. We examined influence of a light dark regime on the spatial association between *B. minutum* and bacteria.
- We employed a novel approach using combination of fluorescence lifetime imaging microscopy with fluorescence *in situ* hybridisation approach to clearly distinguish labelled intracellular bacteria from broad spectrum (450–800 nm) background autofluorescence of *B. minutum*.
- Bacteria were observed inside, tethered to and burrowing into the cell exterior, and at the furrow of dividing cells in *B. minutum*. Significant changes in the abundance of intracellular bacteria relative to autofluorescence in *B. minutum* cells were observed at initiation of light and dark conditions.
- We suggest that the onset of bacterial endosymbiosis is linked to the photoperiod driven changes in *B. minutum* life stages. The re-organisation of thecal plates during cell division of *B.minutum* in dark is likely to result in internalisation of bacteria.

## Introduction

Both free-living and endosymbiotic Symbiodiniaceae, a family of marine dinoflagellate algae, form close associations with taxonomically diverse bacteria and harbour them intracellularly (<u>Lawson *et al.*, 2018</u>; <u>Maire *et al.*, 2021</u>). Possible roles of these bacteria include cycling of sulfur rich compounds (<u>Kiene *et al.*, 2000</u>; <u>Raina *et al.*, 2010</u>; <u>Gao *et al.*, 2020</u>), nutrient cycling and exchange (<u>Matthews *et al.*, 2020</u>), provision of nutrients under nitrogen deplete conditions (<u>Jeong *et al.*, 2012</u>), diazotroph-mediated nitrogen fixation (<u>Rädecker *et al.*, 2022</u>) and induction of *ex hospite* symbiolite formation (aragonite rich calcifying biofilms) (<u>Frommlet *et al.*, 2018</u>; <u>Nitschke *et al.*, 2020</u>). Communities of bacteria residing within the phycosphere (the zone immediately surrounding a cell) of Symbiodiniaceae (<u>Garrido, Amana Guedes *et al.*, 2021</u>) may aid in algal thermal tolerance by provisioning antioxidant pigments (zeaxanthin) to the host (<u>Motone *et al.*, 2020</u>).

The first report of bacteria inside Symbiodiniaceae used 3-dimensional confocal laser scanning microscopy (CLSM) combined with 16S rRNA-targeting fluorescence in situ hybridisation (FISH) (<u>Maire *et al.*, 2021</u>). Diverse bacterial species were found in association with multiple genera of Symbiodiniaceae, and only the intracellular bacteria were highly conserved across Symbiodiniaceae species, suggesting a role in Symbiodiniaceae physiology (<u>Maire *et al.*, 2021</u>). Rod-shaped (perpendicular to) and spiral-shaped bacteria (longitudinally docked) were observed on the cell surface of *Gerakladium* sp. using scanning electron microscopy, and bacteria were observed as randomly distributed in the cytoplasm of *B. minutum* cells by the FISH-CLSM method (<u>Maire *et al.*, 2021</u>).

High background autofluorescence of Symbiodiniaceae makes it difficult to distinguish their associated bacteria when using FISH-CLSM. The broad spectrum autofluorescence (em 450–650 nm) of Symbiodiniaceae cells stems from a range of pigments such as chlorophyll *a* (chl *a*), chlorophyll *c*_2_ (chl *c*_2_), beta-carotene (BC), peridinin (PD), diatoxanthin (Dtx) and diadinoxanthin (Ddx) (<u>Kato *et al.*, 2020</u>; <u>Anthony *et al.*, 2022</u>). Recently, a flavin-based fluorescent protein in freshly isolated Symbiodiniaceae cells from jellyfish, *Cassiopea* sp., was hypothesised to contribute to relatively low levels of autofluorescence in the em 500–600 nm region, compared to BC and Dtx (<u>Anthony *et al.*,</u> <u>2022</u>). Varying levels of background autofluorescence have been reported in unfixed, formaldehyde-fixed and chlorophyll-depleted cells of the dinoflagellate, *Gymnodinium catenatum* (<u>Tang & Dobbs, 2007</u>).

Although several physicochemical methods (<u>Webster *et al.*, 2001</u>; <u>Yakubovskaya *et al.*, 2019</u>; <u>Astafyeva *et al.*, 2022a</u>; <u>Maire *et al.*, 2022</u>) can partially mitigate background autofluorescence, we previously outlined the use of fluorescence lifetime imaging microscopy (FLIM) to study microbial symbiosis (<u>Deore *et al.*, 2022</u>). FLIM measures the fluorescence lifetime of fluorophores rather than their intensity. The fluorescence lifetime of a fluorophore is defined as the average time spent in an excited state before emission of a photon (fluorescence) and return to the ground state (<u>Periasamy & Clegg, 2009</u>). A shorter characteristic autofluorescence lifetime than that of the fluorophores routinely used in FISH is reported in *B. minutum* (<u>Deore *et al.*, 2022</u>).

<u>Maire *et al.* (2021)</u> found that cell cycling and sampling time influenced the prevalence of bacterial present inside the Symbiodiniaceae, *Cladocopium proliferum*. The light-driven cell stage transitions between coccoid and motile forms as well as dividing cells involves a shedding of the cell wall and re-organisation of thecal plates (<u>Wakefield *et al.*, 2000</u>; <u>Kwok *et al.*, 2023</u>). We hypothesised that the surface-attached bacteria may enter *B. minutum* cells while they are undergoing cell wall re-organisation. In our current study, we used FLIM-FISH and evaluated photoperiod driven changes in the relative abundance of bacteria and their detailed intra- and extracellular localisation in the Symbiodiniaceae, *Breviolum minutum* (referred to as the host of intracellular bacteria in this work).

## Materials and methods

### Experimental design and culturing of *B. minutum*

*B. minutum* cells (SCF127-01, Australian Institute of Marine Science, Townsville, Australia) were cultured in Daigo’s IMK (1% w/v, NovaChem, Heidelberg, Australia) medium prepared in filtered Red Sea ^TM^ salt water (fRSS, Reefs Secrets, Burleigh, Australia) and incubated at 26°C under a 12 h:12 h light:dark cycle with 40 µmole m^-2^ s^-1^ light intensity (model no: 740FHC LED light chambers, Taiwan Hipoint Corporation, Kaohsiung, Taiwan) during the light phase.

An exponential phase culture of *B. minutum* of 10^6^ cells mL^-1^ was used. Samples (2 mL) were harvested at eight pre-determined light and dark times over 24 h (Fig. 1). For microscopy purposes, a 1 mL aliquot was centrifuged at 3000 x *g* for 5 min and the pellet was washed with phosphate buffered saline (PBS, pH 7.4). Cells were fixed using 4% paraformaldehyde (PFA, Emgird, Para Hills West, Australia) and stored in 100% ethanol (- 20°C) until FISH was performed. The remaining 1 mL of sample was used to obtain *B. minutum* cells and their intracellular bacteria (ICB) for 16S rRNA gene metabarcoding. Any extracellular and surface-attached bacteria (SAB) were removed as described by <u>Maire *et al*. (2021)</u>. Briefly, samples were filtered using mesh strainers (40 µm; PluriSelect, Leipzig, Germany) and centrifuged (12000 x *g* for 5 min) to retain *B. minutum* cells. The mesh strainers were washed using 500 µL of fRSS. Washed fRSS fractions (stored at -20°C) and unwashed culture (1 mL) of *B. minutum* were analysed by scanning electron microscopy (SEM). Strainers containing fRSS-washed cells were further washed using 500 µL of freshly prepared 6% sodium hypochlorite (Reagent Grade, Sigma) to remove SAB. Washed strainers were stored in sterile microcentrifuge tubes at -20°C until DNA extraction.

**Fig. 1.**
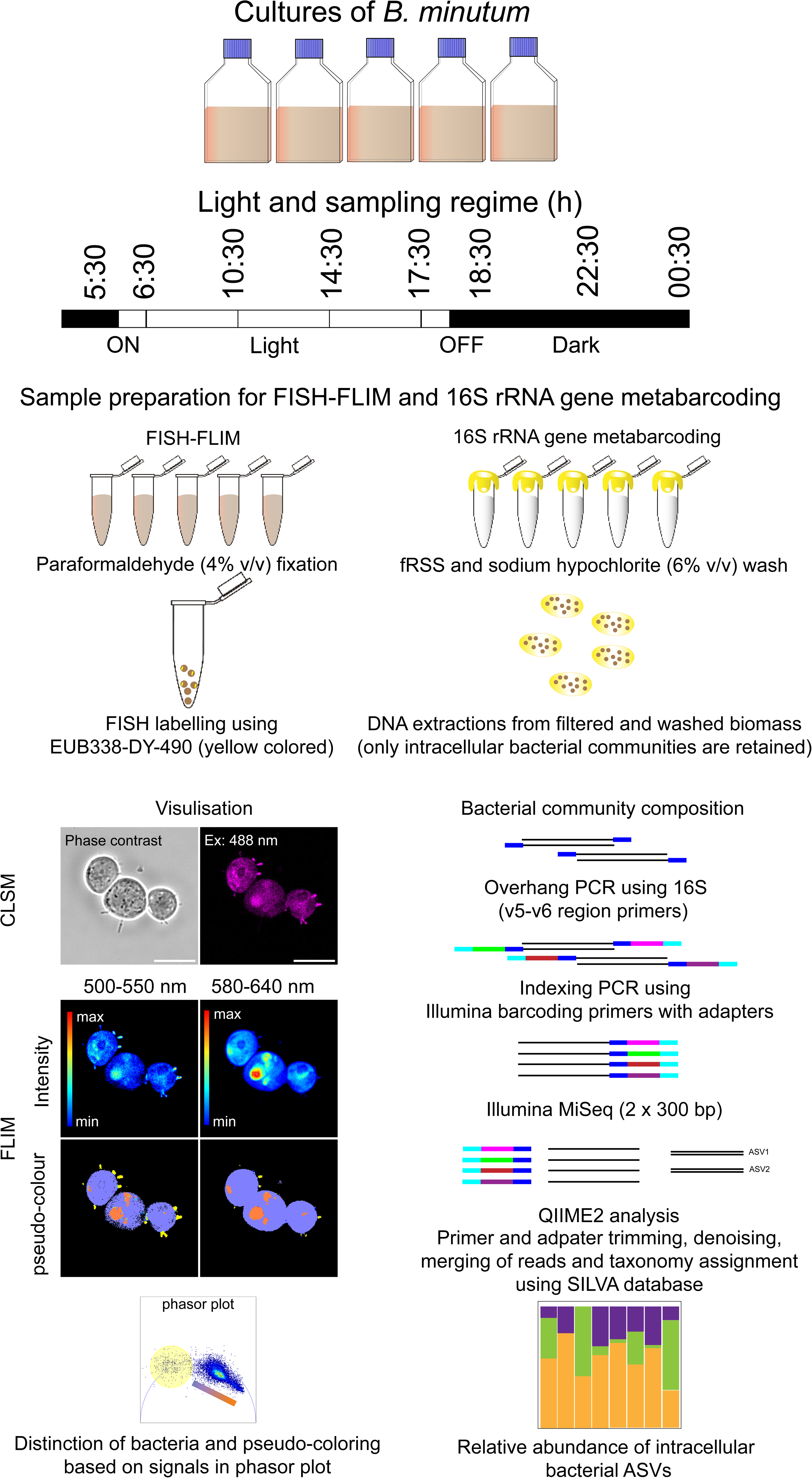
Outline of an experimental design, sampling regime and flow-chart of methods for FISH-FLIM visualisation and 16S rRNA gene metabarcoding for taxonomic identification of intracellular bacteria.

## Microscopy

### Autofluorescence spectra of unfixed and fixed *B. minutum* cells

PFA fixed samples were centrifuged (3000 x *g* for 5 min) and cell pellets were incubated at 40°C to evaporate any remaining traces of ethanol. Dried cells were resuspended in 20 µL PBS mounted on slides, and their autofluorescence spectra were acquired using a CLSM (Nikon A1R) with excitation (ex) lasers at 409, 488, 561 nm and an emission (em) range of 430–650 nm. *B. minutum* cells were mounted on poly-L-lysine (0.1% v/v) coated slides and also observed by CLSM.

### Fluorescence *in situ* hybridisation

Fixed samples were centrifuged at 3000 x *g* for 5 min and cell pellets were incubated at 40°C for 5 mins. Cells were resuspended in 80 µL of hybridisation buffer (HB) containing 20 mM of Tris-HCl (pH - 7.4), 0.9 M NaCl, 0.01% sodium dodecyl sulfate (SDS) and 25% formamide. For 16S rRNA labelling, the EUB338 probe with the DY-490 fluorophore (EUB338-DY490; ex and em_maximum_ 491 and 515 nm, respectively; Biomers, Ulm, Germany) was added at 2.5 ng µL^-1^. Samples were incubated at 46°C in dark for 3-4 h with occasional mixing. Samples with non-EUB338 probe containing the ATTO-550 fluorophore (ex and em_maximum_ 554 and 575 nm, respectively; Biomers, Ulm, Germany) served as a negative control. Samples were centrifuged at 3000 x *g* for 5 min and the HB removed by pipetting. Unbound probe was removed by resuspending cells in 500 µL of pre-warmed (48°C) wash buffer (WB) containing 20 mM of Tris-HCl (pH 7.4), 0.149 M NaCl, 0.01% SDS and 40 mM EDTA, centrifuging (3000 x *g* for 5 min) and pipetting off the supernatant. Cells were resuspended in WB and incubated at 48°C in dark for 20 min followed by centrifugation at 3000 x *g* for 5 min. The cell pellet was resuspended in 50 µL of PBS and stored in -20^°^C. For FLIM image acquisition, 10 µL of FISH labelled samples were placed onto glass slides, covered by #1.5 coverslips.

### Confocal Laser Scanning Microscopy coupled with Fluorescence lifetime imaging microscopy

The bacteria labelled using EUB338-DY-490 were visualised using an Olympus FV3000 CLSM coupled to a 488 nm pulsed laser operated at 80 MHz and an ISS A320 FastFLIM box (ISS Medical DBA ISS Inc., Champaign, USA). The 488 nm pulsed laser maximally excited DY-490 while also generating broad spectrum autofluorescence from *B. minutum* and the FastFLIM box enabled time resolved distinction of these two fluorescent signals. The collective fluorescence from DY-490 labelled *B. minutum* was directed through a 405/488/561 dichroic mirror and then split by a 550 nm long filter between two external photomultiplier detectors (H7422P-40 of Hamamatsu Corporation, Japan) fitted with 520/50 nm (CH1-green) and 600/50 nm (CH2-red) bandwidth filters. This detection scheme resulted in the fluorescence from DY-490 (em 515±20 nm) being maximally captured in CH1 while *B. minutum* autofluorescence was detected in both channels. The signal detected in CH1 and CH2 was processed by the ISS A320 FastFLIM box data acquisition card to report the phasor coordinates of the fluorescence lifetime associated with each pixel (FLIM data). All two channel FLIM data were acquired with a 60x water immersion objective 1.2 numerical aperture within a 212 x 212 µm field of view (FOV) containing 1–10 *B. minutum* cells using a 256-pixel frame size, 20 µs pixel dwell time. For each replicate sample, two channel FLIM data were acquired in four different FOVs using the ISS Vista Vision software and a pre-calibration of the instrument as well as phasor space against fluorescein, which at pH 9 has a single exponential lifetime of 4.04 ns. Samples from all experimental time points were randomised to eliminate bias associated with variation in laser intensities and instrument responses on a given day.

### FLIM image processing

All FLIM data were analysed in SimFCS4 (software developed by the Laboratory for Fluorescence Dynamics, University of California, USA) by the phasor approach to fluorescence lifetime analysis that is a well-established fit-free method for quantification of multi-component FLIM data (<u>Digman *et al*., 2008</u>; <u>Hinde *et al*., 2012</u>; <u>Ranjit *et al*., 2018</u>). In brief, the phasor approach transforms the fluorescence lifetime recorded in each pixel of a FLIM image into a vector (a phasor), which when represented in a two-dimensional coordinate system (a phasor plot) enables: (1) the fractional contribution of two or more independent molecular species coexisting in the same pixel (e.g., DY-490 and autofluorescence) to be determined by vector algebra, and (2) spatial mapping of these different molecular species identified throughout source FLIM images. To enable these analytical capabilities in the context of the two channel FLIM data presented in supplementary sheet (Fig. S1), the phasor coordinates of pure DY-490 and *B. minutum* autofluorescence were first independently determined by FLIM acquisition of a pure culture of *Marinobacter adhaerens* labelled with EUB338-DY-490 and a sample of unstained *B. minutum.* The fluorophore, DY-490 exhibit a single exponential fluorescence lifetime of 2.9 ns (s, g are 0.571, 0.236) on the line of universal circle (yellow cursor; Figs. S1a–d and e) in pure FISH labelled *M. adhaerens* samples. The lifetime of DY-490 is shifted to 3.1 ns (s, g are 0.456, 0.289) in samples of mix *B. minutum* and labelled bacteria (Figs. 2c-f). *B. minutum* autofluorescence exhibit a linear combination of multiexponential fluorescence lifetimes (blue orange coloured palette gradient bar; Figs. S1e). These phasor coordinates were then employed to define a phasor-based palette that could pseudo-colour the position of DY-490 labelled bacteria within *B. minutum* samples and then quantify of the fraction of pixels positive for DY-490 labelled bacteria inside of *B. minutum* cells. Pseudo-coloured FLIM maps prior to quantification were subjected to a spatial median filter and a minimum intensity threshold to remove phasors calculated from pixels with insufficient counts for phasor analysis. These cursor assignments were later used for pseudo-colouring of images indicating the DY-490 signal (yellow coloured) within autofluorescent (blue orange coloured palette gradient bar) *B. minutum* cells (Fig. S1). The percentages of DY-490 signal inside *B. minutum* cells were determined by creation of an ellipse-shaped mask encompassing the total cell area. The feature counts of DY-490 in CH1 within the masked area (*B. minutum* cell) were normalised by autofluorescence counts in CH1 or CH2 (the channel with highest signal counts was considered on each occasion). Counts of DY-490 in each FOV were further divided by the number of *B. minutum* cells present in the FOV and expressed as % relative DY-490 counts per Symbiodiniaceae cell. The mean values from 5 FOVs of each biological replicate (n = 5) were averaged for each experimental time point. Based on the position of bacteria (inside or surface-attached), *B. minutum* cells were allocated into three categories: intracellular ICB, SAB or no bacteria (NB). The percentage of *B. minutum* cells harbouring ICB, SAB and NB were visually estimated for each FOV. Cells containing both ICB and SAB were categorised once in the ICB category. CLSM images were processed in Fiji (1.53t, NIH, USA) to separate colour channels and assign scale bars. All microscopy images were compiled in combination of the figures in Adobe illustrator, version 26.3.1 (USA).

**Fig. 2.**
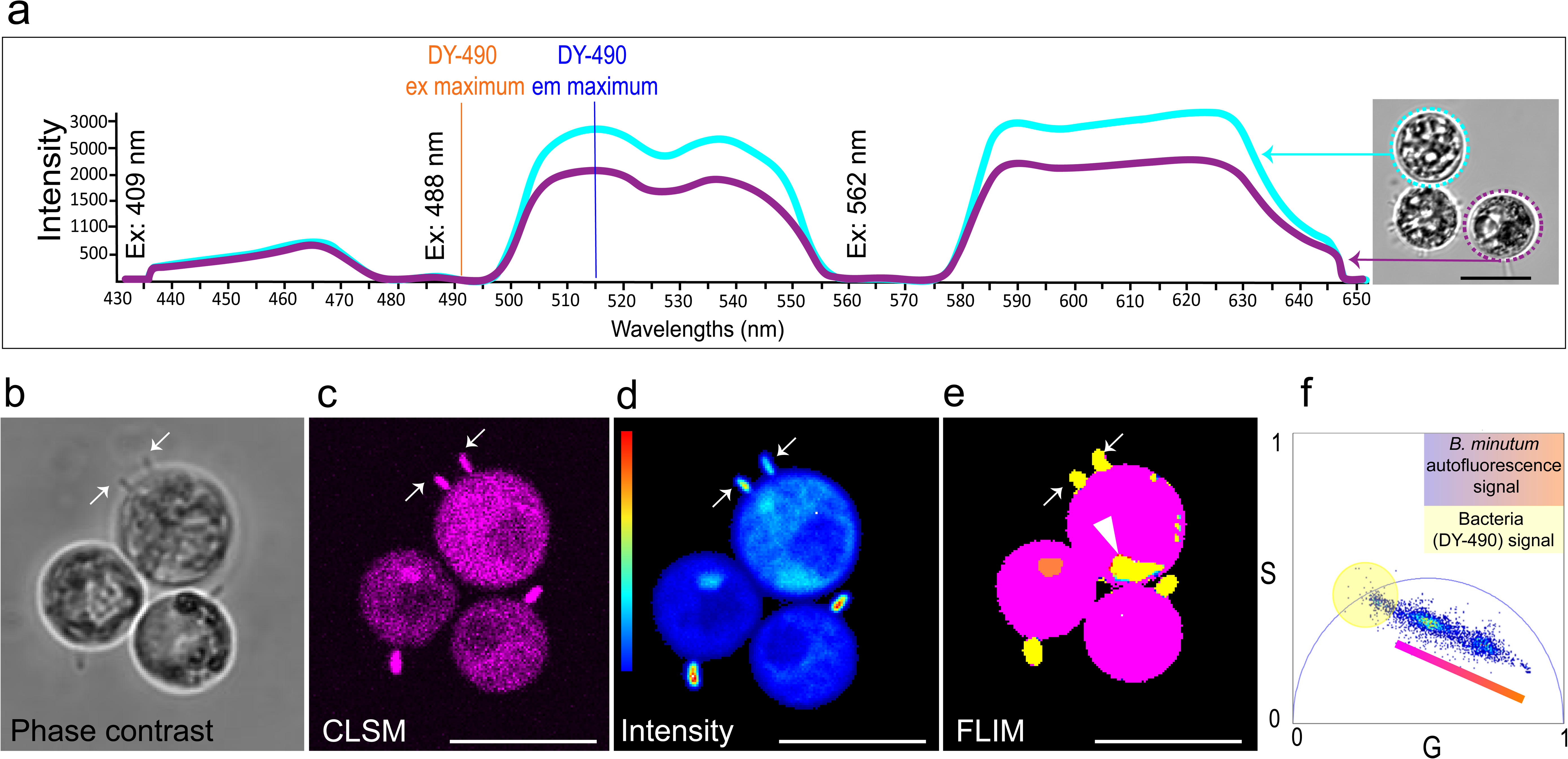
Background autofluorescence in unstained *B. minutum* and distinction of bacteria labelled with EUB338-DY-490 probe targeting 16S rRNA using fluorescence lifetime image microscopy (FLIM) a) Broad spectrum autofluorescence by confocal laser scanning microscopy (CLSM) (ex 409, 488 and 561 nm, cyan and magenta lines corresponding to *B. minutum* cells shown in dotted circles of similar colour). Vertical dotted lines indicate ex _maximum_ and em_maximum_ (491 and 515 nm, respectively) of DY-490 which overlaps with *B. minutum* autofluorescence. b–e) All show three *B. minutum* cells in the same field of view and white arrows indicate surface-attached bacteria. b) Phase contrast image showing surface-attached rod-shaped bacteria c) CLSM image (ex 488 nm; em 500-550 nm) of *B. minutum* cells with surface attached bacteria d) Intensity (in the colour scale on the left, blue is minimum and red is maximum value of emission range 500–550 nm), e) FLIM image pseudo-coloured using colour palette defined in Fig. f indicating the presence of intracellular bacteria (yellow coloured and white arrowhead) that were indistinguishable in Fig. c. f) The phasor plot showing DY-490 and *B. minutum* specific autofluorescence lifetimes in yellow coloured cursor and pink orange coloured palette gradient bar, respectively. Scale bars = 10 µm.

### Scanning electron microscopy

*B. minutum* cells in the fRSS-washed fractions were obtained from -20^°^C stored filters and resuspended in filter sterilised IMK. Both washed and unwashed samples were centrifuged at 5000 x *g* for 5 min and pellets were resuspended in 200 µL of filter sterilised IMK containing 2.5% (v/v) of glutaraldehyde. Samples were incubated at (room temperature) RT for 60 min and were subjected to serial washes (5000 x *g* for 5 min) using 500 µL fRSS; 250 µL of fRSS and PBS; 500 µL of PBS, and two subsequent washes with 500 µL of MilliQ water. Washed samples were placed on poly-L-lysine (0.1% v/v) coated coverslips and incubated at RT for 30 min to allow *B. minutum* cells to adhere. Cells adhered to coverslips were serially dehydrated once in 10%, 20%, 40%, 60%, 80%, and thrice in 100% ethanol for 30 min each. Dehydrated samples were stored in 100% ethanol at 4^°^C until critical point drying was performed in a Leica CPD30 for 3 h. Samples were coated with gold using a Quorum Q150T ES plus for 2 min prior to observation under a XL30 scanning electron microscope (Philips) at 5 kV beam (BioSciences Microscopy Unit, University of Melbourne).

### Metabarcoding (16S rRNA gene) of intracellular bacterial communities

DNA was extracted from biomass on filters using a method outlined by <u>Wilson *et al.* (2002).</u> Briefly, 0.375 mL of freshly prepared filter sterilised (0.22 µm) extraction buffer (Tris buffer pH-9, 100 mM EDTA, 100 mM NaCl; 1% SDS, 0.2 mg mL lysozyme) was added to a 1.5 mL microcentrifuge tube with the biomass-containing filters. Tubes were vortexed and incubated at 37°C for 30 min followed by Proteinase K (0.5 mg mL) treatment. Samples were lysed using ∼100 mg sterile glass beads (425–600 µm diameter) followed by bead beating for 30 s at 30 Hz (Qiagen tissue-lyser II). Samples were incubated at 65°C for 60 min. DNA was precipitated using 0.45 mM ice-cold potassium acetate followed by 30 min of incubation on ice. Samples were centrifuged twice at 15000 x *g* for 5 min and the supernatant was treated with RNAse (0.13 mg mL) at 37°C for 30 min. DNA was precipitated using 500 µL of ice-cold iso-propanol and incubated at RT for 15 min. DNA was pelleted by centrifuging samples at 15000 x *g* for 15 min and washed twice with ice-cold ethanol. Samples were air dried at RT and resuspended in MilliQ water.

The ICB community composition was characterised in bulk Symbiodiniaceae cells using 16S rRNA gene metabarcoding. Sample were amplified using 16S rRNA gene primers spanning hypervariable V5 and V6 regions. The amplified products were used for library preparation followed by indexing PCR (see supplementary sheet S1 for primer oligonucleotide sequence, detailed PCR and library preparation conditions). Samples were sequenced on a single Illumina MiSeq run using v3 (2 x 300 bp) at WEHI.

### Amplicon sequence variant (ASV) analysis

The raw sequence reads were processed in QIIME 2.2022-8 (<u>Bolyen *et al*., 2019</u>) for trimming of the Illumina adaptors using cutadapt plugin (<u>Martin, 2011</u>). Quality plot visualisation, raw read filtering and demultiplexing of trimmed reads were performed using demux plugin (<u>Bolyen *et al*., 2019</u>). The DADA2 plugin was used for truncation of sequences below the medium quality score of Q35, denoising, chimera detection and dereplication of reads prior to assembling paired end reads (<u>Callahan *et al*., 2016</u>). A summary statistic table was generated to ensure successful pre-processing of reads. Taxonomy identifications were assigned to ASVs by training a naive Bayes classifier with the feature classifier plugin (<u>Bokulich *et al*., 2018</u>; <u>Bolyen *et al*., 2019</u>) which classifies ASVs based on a 99% similarity score to the V5 and V6 regions of 16S rRNA genes (784F/1061R primer pair). ASVs from mitochondria and chloroplasts were filtered out. Files containing metadata, phylogeny tree, and tables with ASV taxonomic classifications were imported in R studio for statistical analysis and visualisation.

### Statistical analysis of ASVs and visualisation in R studio

R studio (version 4.1.2) with phlyoseq, ape, microbiome, decontam, and ggplot2 packages were used for data analysis and visualisation. The statistical level of significance was 0.05. ASVs present in the extraction blanks and no DNA template controls were removed using the prevalence method (threshold p = 0.1) in the decontam package of R (<u>Davis *et al*., 2018</u>). Alpha diversity metrics (observed ASVs, Simpson’s and Shannon’s indices) were estimated by rarefaction (i.e., common lowest read number across all samples) of ∼4709 reads per sample and sufficient depth coverage to capture maximal diversity across all samples. Overall differences in alpha diversity were analysed by the non-parametric Kruskal–Wallis test. Differences in the community composition across samples were estimated using weighted unifrac dissimilarity matrices and tested by permutational multivariate analysis of variance (PERMANOVA) across all time points. Differences in the ICB community composition across all experimental time points and DNA template volumes were visualised using principal coordinate analysis (PCoA). The top 15 abundant genera of intracellular bacterial communities across all timepoints were visualised as stacked bar plots in R studio.

### Statistical analysis

The effect of photoperiod (different time points in light and dark) on the type of spatial association of bacteria (ICB, SAB and NB) and *B. minutum* was assessed by a repeated measure mixed-effect model (one way ANOVA). Sidak’s multiple comparison test was used to test differences in *B. minutum* proportions containing ICB compared to SAB and NB across all time points. The difference in relative abundance of ICB (based on DY-490 signals) per *B. minutum* across light and dark time points was assessed by mixed effects analysis and Tukey’s test was used for multiple comparisons. The level of significance for all statistical analyses was 0.05. Data were visualised using Graphpad Prism (version 9.5.0, USA).

## Results

### Distinction of DY490-labelled bacteria from *B. minutum* autofluorescence

Fig. 2a shows the broad spectrum autofluorescence of unfixed *B. minutum* cells (ex lasers 409, 488 and 562 nm; em 430–650 nm) overlapping with the emission of DY-490 (em_maximum_ 515 nm). The autofluorescence intensity of *B. minutum* cells (cyan and magenta-coloured dotted circles, Fig. 2a) was higher in em range 500–555 nm and 580–645 nm compared to 435–475 nm. Similar autofluorescence spectra were observed from PFA fixed *B. minutum* cells although signal intensities were dampened by ∼60% (see supplementary sheet S1, Fig. S2). Surface-attached rod-shaped cells are clearly visible in phase contrast microscopy (white coloured arrow, Fig. 2b) and they coincide with EUB338-DY-490 labelled bacteria (Fig. 2c) in CLSM. The detection of any intracellular bacteria was hindered by background autofluorescence in all three *B. minutum* cells (Fig. 2c). However, FLIM images and analysis using a fit-free phasor approach shows clear resolution of ICB as well as SAB (Figs. 2d and e, white arrowhead and arrows, respectively) from high background autofluorescence (pink orange coloured gradient bar in Fig. 2f, respectively).

### Phasor signature of DY-490 and *B. minutum* autofluorescence

In Fig. S1, cells of *M. adhaerens* (EUB338-DY-490 FISH probe labelled, Fig. S1a, b and e) showed a distinct single exponential lifetime of 2.9 ns (s, g are 0.571, 0.236). The positioning of DY-490 on the phasor plot was shifted to 3.1 ns (s, g are 0.456, 0.289) in mixed samples of bacteria and *B. minutum* (Figs. 2d–f). Regardless the difference in lifetime of DY-490 (labelled bacteria) and *B. minutum* was evident from the unique positioning of multiexponential lifetimes associated with autofluorescence in blue orange coloured palette gradient bar in phasor plot (Fig. S1c–e).

### Localisation of *B. minutum*-associated bacteria

In Fig. 3, bacteria were localised to three different *B. minutum* niches i.e., intracellularly (ICB) (blue arrowhead, Fig. 3f), adjacent to/lying along cell surfaces (SAB) (Figs. 3f, i) and occasionally burrowing into the cell surface (cyan-coloured arrowhead in Fig. 3i), during a diurnal cycle of *B. minutum*. Rod-shaped bacterial cells (cyan-coloured arrows in Figs. 3b, c, e, f, h, and i) were frequently attached to the cell surface of *B. minutum* cells throughout the light and dark periods. SEM images (Figs. 3j-m) show that short rod-shaped (cyan-coloured arrows) and long spiral-shaped (blue-coloured arrows) microbial cells were attached to the cell surface of unwashed and fRSS-washed *B. minutum* cells. Unwashed cells (Fig. 3j) had much surrounding material likely to be microbial cells and organic matter. *B. minutum* cells also contained ICB (blue arrowhead in Fig. f) surrounding the pyrenoid (orange coloured arrow in Figs. 3d and e) where levels of autofluorescence were slightly lower compared to the surrounding cellular matrix (CLSM, Fig e). The fluorescence signals from ICB and burrowing bacteria in CLSM images (Figs. 3e and h) were masked because of *B. minutum* autofluorescence, especially for the burrowing rod-shaped bacterium as it is beside a bright distinct orange coloured body (Fig. 3i) which emits strong autofluorescence (Fig. 3h).

**Fig. 3.**
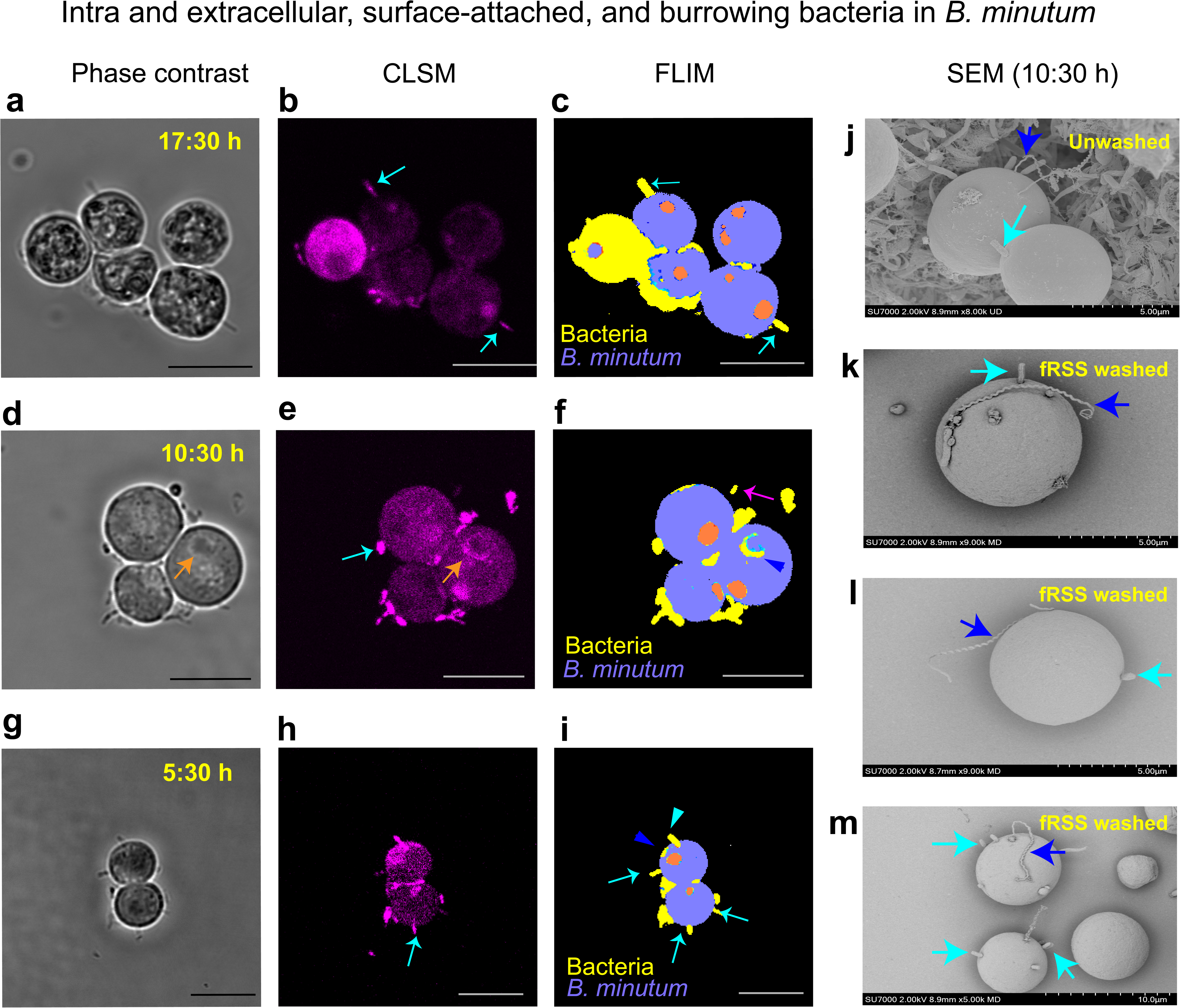
Localisation of bacteria (surface attached, intracellular, burrowing and extracellular) in *Breviolum minutum* cells in different time points of the diurnal cycle using combined fluorescence *in situ* hybridisation (FISH) and fluorescence lifetime imaging microscopy (FLIM), and scanning electron microscopy (SEM). The same fields of view of cells at a–c) 17:30 h (light), d–f) 10:30 h (light), g–I) 5:30 h (dark). Images in Figs. a, d, g are phase contrast images; Figs. b, e, h are confocal laser scanning microscopy (CLSM); Figs. c, f, I are pseudo coloured FLIM images. In Fig. d, the orange arrow indicates the pyrenoid. In Fig. f, EUB338-DY-490 labelled bacteria (coloured yellow) were found intracellularly (blue arrowhead) and extracellularly (pink arrow) (FLIM images) in *B. minutum*. In Fig. i, a rod shaped bacterium burrowing into the surface of *B. minutum* (cyan arrowhead) is shown and in Figs. b–i, surface-attached (cyan arrows) bacteria are shown. CLSM and FLIM images are presented to highlight challenges in distinction of fluorescently labelled bacteria (surface-attached vs. burrowing and intracellular bacteria) from *B. minutum* autofluorescence (ex 488 nm, em 500–550 nm). j–m) SEM of closely associated surface-attached rod and spiral-shaped bacteria (cyan and blue-coloured arrows, respectively) to *B. minutum* cell wall. Scale bars = 10 µm.

### Percent of *B. minutum* cells associated with bacteria

The percentages of *B. minutum* cells (Fig. 4) with ICB, SAB or NB were largely influenced by the time of the photoperiod (p = 0.025, F [14, 84] = 2.02). Over a 24 h photoperiod, the percentage of *B. minutum* cells with NB (40–65%) were significantly higher compared to *B. minutum* cells with ICB (5–22%) across all time points (5:30 h, p = 0.0001; 6:30 h, p<0.0001; 10:30 h, p<0.0001; 17:30 h, p = 0.0013; 18:30 h, p = 0.027; 00:30 h, p = 0.0009) except for 14:30 h and 22:30 h. At pre-dawn (5:30 h) and pre-dusk (17:30 h), only 5.9% (p = 0.041) and 6.6% (p = 0.099), respectively, of *B. minutum* cells had ICB. The number of *B. minutum* cells with SAB increased during the light phase (6:30–17:30 h) while cells with ICB remained stable. There were no significant differences in the percentage of *B. minutum* cells with SAB in the light and dark phases (see supplementary sheet S1, Fig. S3). At the following times after dusk, 18:30 h, 22:30 h and 00.30 h, 15%, 22% and 20%, respectively, of *B. minutum* cells had ICB (Fig. 4).

**Fig. 4.**
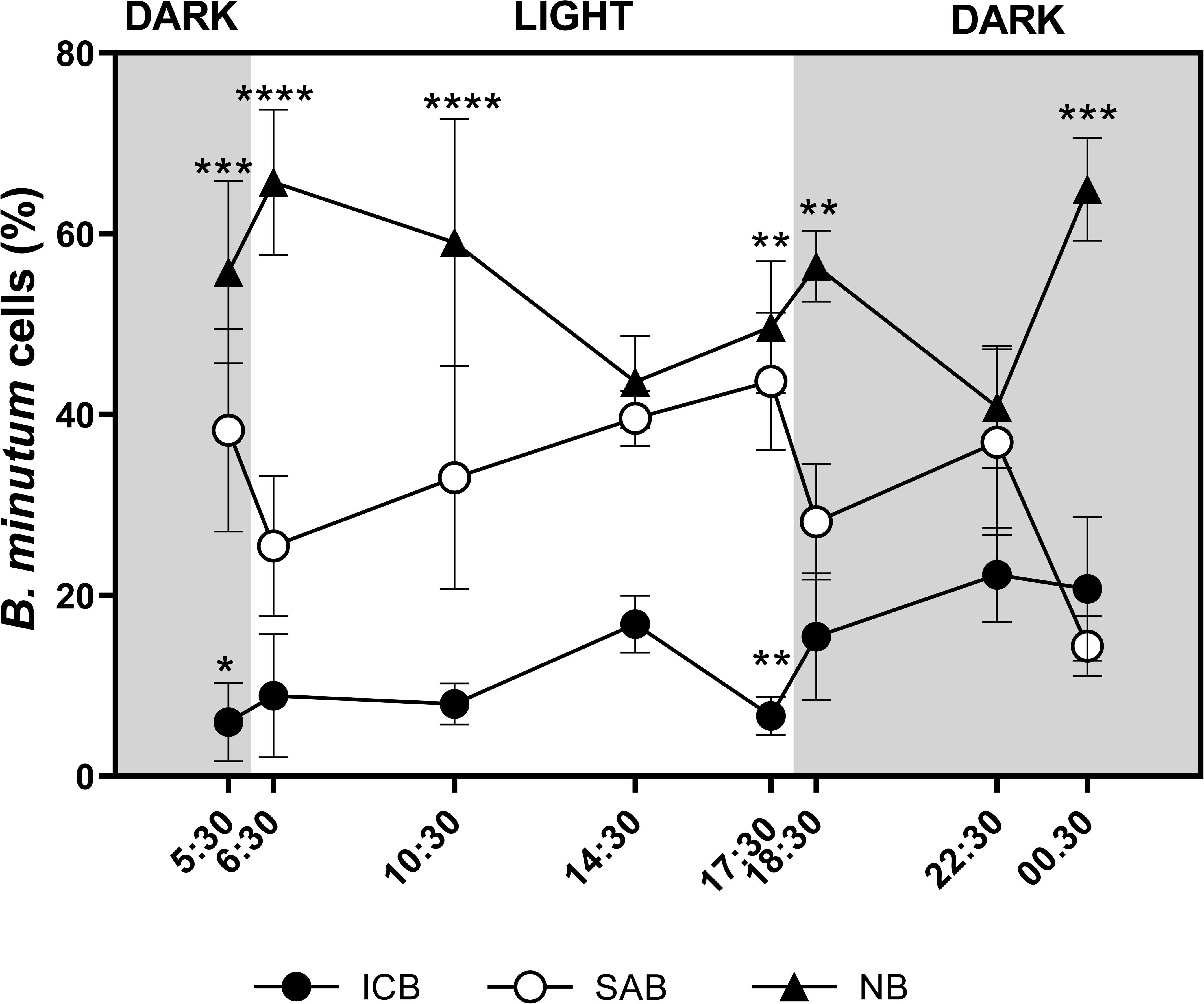
Percentage of *Breviolum minutum* cells containing intracellular (ICB), surface-attached (SAB) bacteria and no bacteria (NB) during the light (6:00 h to 17:30 h) and dark (18:30 h to 5:30 h) phases. Data points are percent mean cell numbers of *B. minutum* across all biological replicates (n = 5) and estimate of errors are the standard error of the mean. p-values are *0.041, **0.009, ***0.0001 and ****<0.0001 where the cut-off for significance was 0.05. Shaded grey and white areas in the plot indicate dark and light phases of a 24 h photoperiod, respectively.

### Relative abundance of ICB associated with *B. minutum*

The counts of DY-490 signals from ICB relative to *B. minutum* autofluorescence signals significantly varied over a 24 h photoperiod (mixed model ANOVA, p = 0.0019, F [4, 15] = 2.905) (Fig. 5). The relative abundance of ICB in *B. minutum* cells at 18:30 h (30 min after dusk) was on average 5 (p = 0.0083) and 9.5-fold (p = 0.0082) higher compared to their abundances at 5:30 h and 6:30 h (30 min pre- and post dawn, respectively) (Fig. 5). During the dark phase (18:30 h to 5.30 h), ICB were 2.5-fold more abundant (p = 0.0315, Welch’s t-test t = 2.293 and df = 22.69) than they were in the light phase (see supplementary sheet S1, Fig. S4). No correlation was observed between relative abundance of ICB (normalised DY-490 feature counts) and percentage of *B. minutum* cells that harboured bacteria, meaning that when more *B. minutum* cells harboured ICB, the abundance of ICB per *B. minutum* cell did not increase (see supplementary sheet S1, Fig. S5).

**Fig. 5.**
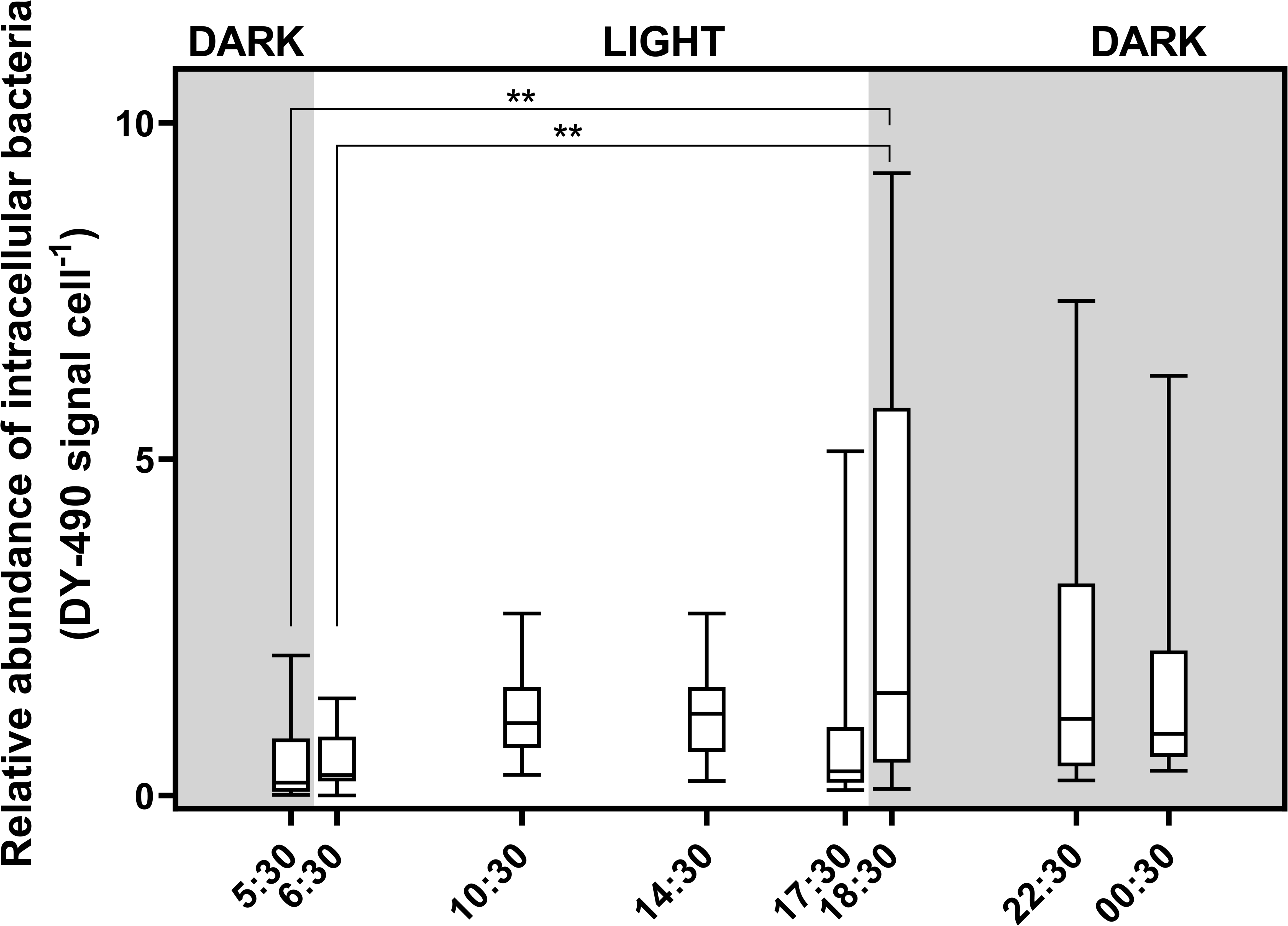
Abundance of intracellular bacteria (ICB) based on feature counts of DY-490 normalised to autofluorescence signal counts per *Breviolum minutum* cell during the light and dark phases (6:00–17:30 and 18:30–5:30 h, respectively). Data are mean of independent biological replicates±SD (n = 5) and horizontal lines within each bar indicates median values. Asterisks (**) indicate p-value of 0.008 where the cut off for significance was 0.05.

### Presence of ICB in coccoid and dividing *B. minutum* cells

Symbiodiniaceae go through cell cycles regulated by light and dark phases, where light drives the growing/DNA synthesis (i.e., G1 to S to G2/M) stage and dark is required for mitotic division (or cytokinesis i.e., G2/M to G1) (<u>Wang *et al*., 2008</u>). In coccoid *B. minutum* cells at different times of the photoperiod (pink dashed circled cells in Figs. 6a-d), ICB were randomly distributed throughout cytoplasm, occasionally adjacent to distinct autofluorescent bodies (orange coloured in FLIM images) and alongside the cell membrane (white arrows Fig. 6b). A membranous-like feature surrounded pairs of dividing *B. minutum* cells (Fig. 6e and inset phase contrast images in Fig. 6i-l) and bacteria were observed at the site of a furrow or invagination (furrow, Fig. 6e, and bacteria, Figs. 6h-l, is indicated by blue and cyan arrowheads, respectively) of early stage dividing *B. minutum* cells (clarified in phase contrast inset images, Figs. 6h-i). Dividing cells of *B. minutum* frequently contained two and occasionally three distinct autofluorescent bodies (orange coloured in FLIM images, Fig. 6h, k, l). Occasionally these bodies were absent in late stages of cell division (Figs. 6i, j). However, they were distributed throughout the *B. minutum* cytoplasm in earlier stages of cell division when the division septum was visible but invagination had not yet commenced (Figs. 6k, l).

**Fig. 6.**
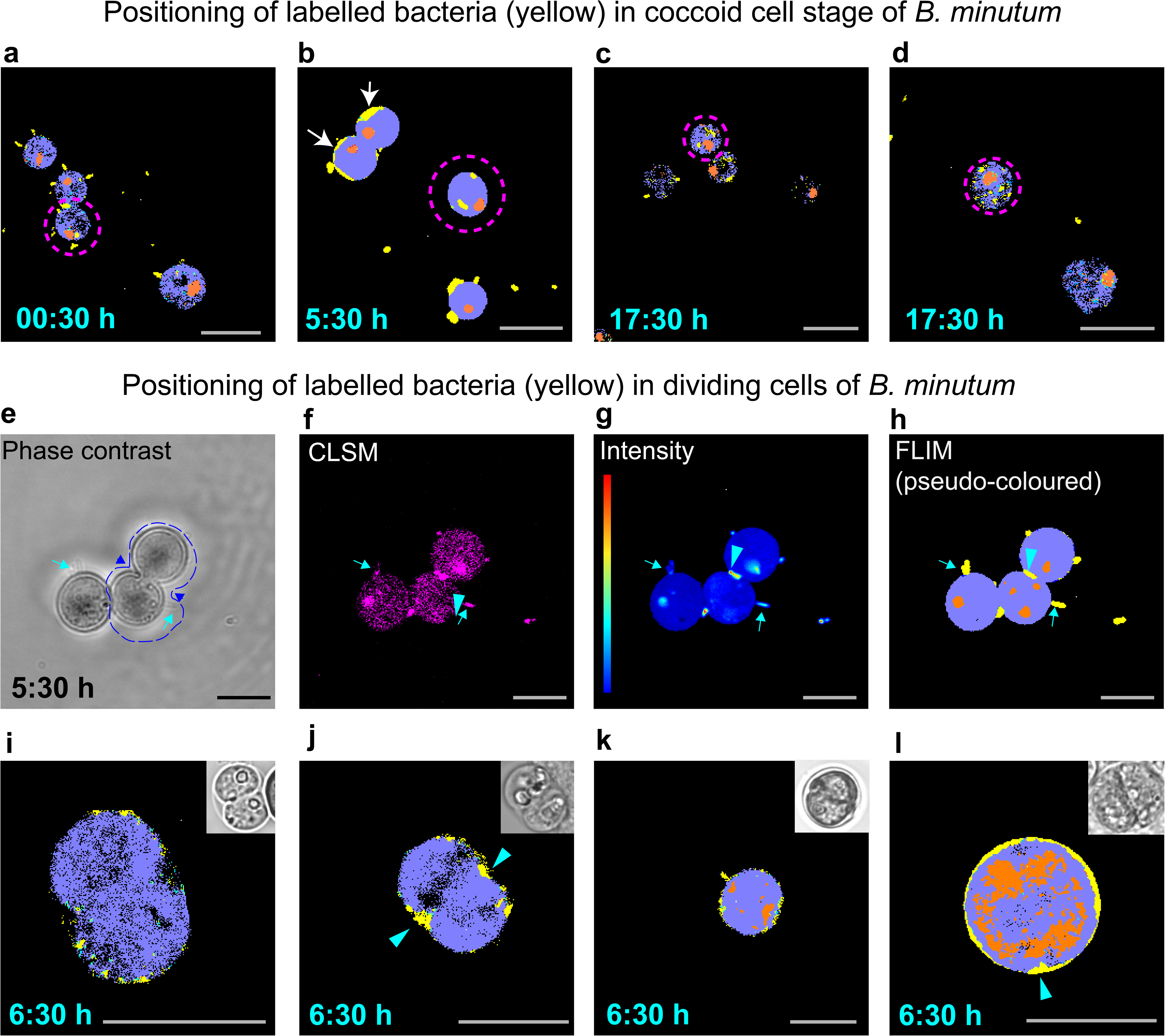
Position of EUB339-DY-490 labelled intracellular (ICB) and surface attached (SAB) bacteria (yellow coloured) in coccoid and dividing *Breviolum minutum* cells. a–d) In pink dotted circled *B. minutum* cells, ICB are adjacent to a distinct autofluorescent body (orange coloured). In Fig. b) white arrows show ICB alongside of the *B. minutum* cell membrane. e– h) The same field of view of one coccoid and one dividing *B. minutum* cells. SAB on *B. minutum* cells is indicated by cyan arrows. i–l) Different *B. minutum* cells in dividing stage. In Figs. e–l), presence of bacteria (cyan arrowheads) with no distinct cell shape at a furrow (blue arrowheads, site of cleavage) of a dividing *B. minutum* cell in phase contrast Fig e, and in a sheath-like layer surrounding the cell membrane (visible in inset phase contrast images in Figs. i–l). Scale bar = 10 µm.

### Intracellular bacterial taxonomic affiliation (16S rRNA gene metabarcoding)

A total of 7305947 raw reads were obtained from the Illumina MiSeq sequencing of ICB obtained from sodium hypochlorite-washed *B. minutum* cells. Data cleaning steps such as merging of paired end reads, denoising and chimera removal resulted in a dataset of 4185177 reads. After removal of contaminant ASVs (identified in no template controls and extraction blanks), 6122 ASVs remained. No DNA template concentration related variation in ASVs was observed across all time points (see supplementary sheet S1 and S2, Figs. S6 and 7). The top 15 ASVs across all timepoints were from genera - *Brachybacterium*, *Sphingomonas*, *Cutibacterium*, *Cognatishimia*, *Massilia*, *Marinobacter*, *Endozoicomonas*, *Staphylococcus*, *Methylobacterium*, *Bacterioides*; unknown genera from the families Moraxellaceae, Flavobacteriaceae, Rhodobacteraceae, order Micrococcales and an uncultured species (AB1) belonging to family Rickettsiaceae (Fig. 7a). There was no relationship between sampling time (in light or dark) and ICB community structure according to the PCoA (Fig. 7b). Nor was there a statistically significant variation in the ICB composition over time (PERMANOVA, weighted unifrac method, Df = 7, F-value = 1.1064, p-value = 0.125). The alpha diversity (difference within samples based on observed OTUs) was not significantly different across all timepoints (χ2 = 4.5628, df = 7, p-value = 0.7131, see supplementary sheet S1, Fig. S6). No grouping of samples based on dissimilarity matrices (beta diversity) was observed in samples in relation to photoperiod (see supplementary sheet S1, Fig. S7). The presence of *Cutibacterium* sp. is not unusual as this bacterium is widely reported as a kit contaminant (<u>Hartman, 2020</u>; <u>Maire *et al*., 2021</u>). *Endozoicomonas* sp. is observed for the first time in cultured *B. minutum* and is often dominant in corals where it forms cell-associated microbial aggregates in coral tissues (<u>Maire *et al*., 2022</u>). However, localisation of *Endozoicomonas* sp. in the proximity of *in hospite* Symbiodiniaceae is not consistently observed in corals (<u>Maire *et al*., 2022</u>; <u>Gardner *et al*., 2023</u>). The ASVs associated with *Endozoicomonas* sp. found in this study were a close match with two of eleven pure cultures of *Endozoicomonas* sp. isolates maintained in the lab (data not shown). Therefore, we hypothesise that the presence of *Endozoicomonas* in our *B. minutum* cultures is likely the result of laboratory cross-contamination.

**Fig. 7.**
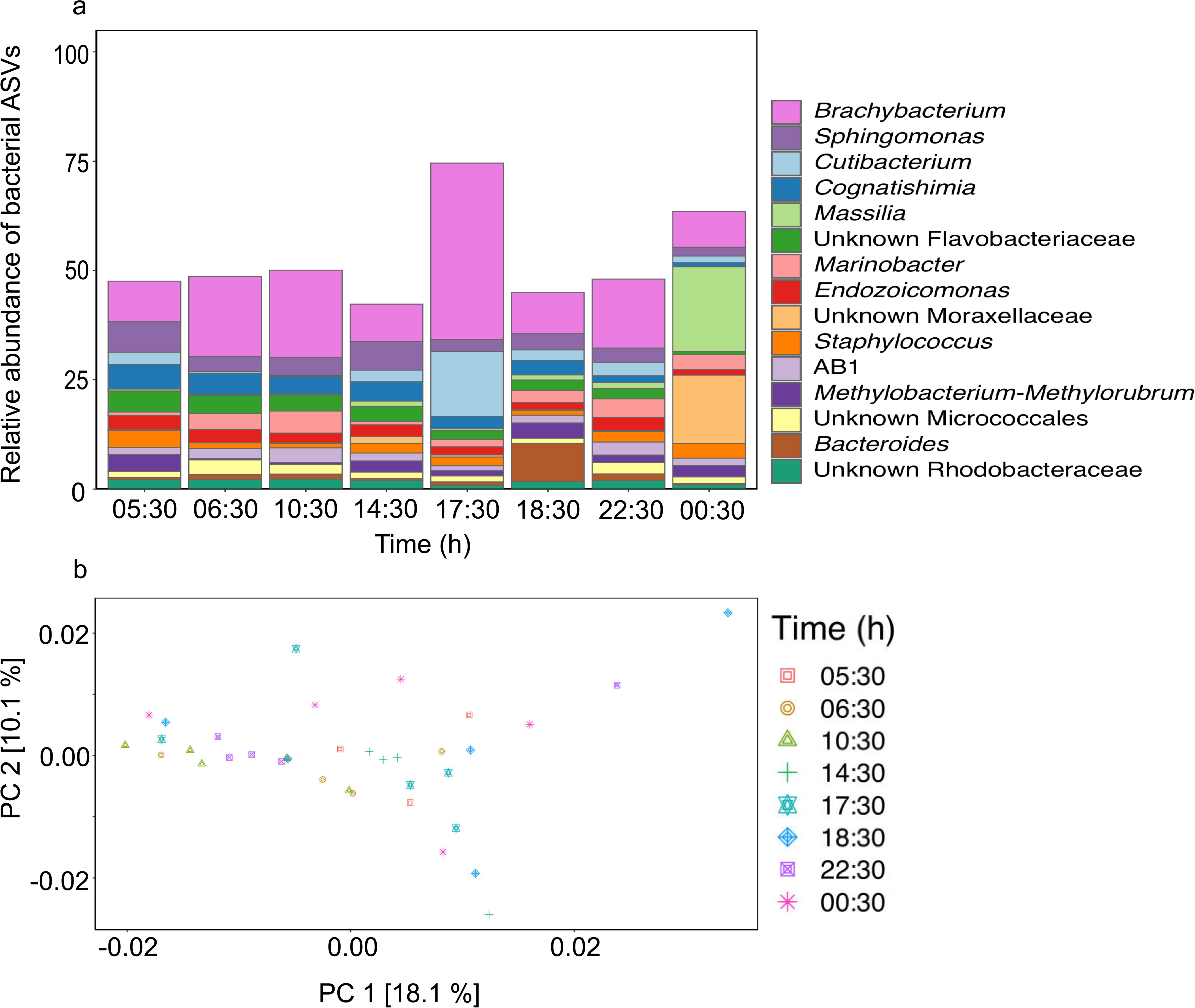
Diversity of intracellular bacteria (ICB) communities in *Breviolum minutum* cells during light (6:00–17:30 h) and dark (18:30–5:30 h) phases. a) Relative abundance of the top 15 bacterial genera as a function of time. b) Principal coordinate analysis (PCoA) for visualization of beta diversity of ICB based on weighted unifrac dissimilarity matrix. Data represents percent distribution of bacterial ASVs averaged from independent biological replicates.

## Discussion

### Autofluorescence in *B. minutum*

The observed levels of autofluorescence in *B. minutum* cells (Fig. 2) are likely to arise from fluorescent pigments chl *a*, chl *c*_2_ and PD which are the major components of the light harvesting complex in Symbiodiniaceae (<u>Jiang *et al*., 2014</u>). High autofluorescence from these pigments hinders the use of fluorescence-based observational methods to study intracellular symbionts of Symbiodiniaceae (<u>Maire *et al*., 2021</u>).

In *B*. *minutum* (strain SSB01), autofluorescence (ex 488 nm, em 501–550 nm) from an accumulation body containing amyloid-like substances (<u>Yonge, 1931</u>; <u>Freudenthal, 1962</u>) was reported by <u>Gornik *et al*. (2022)</u>. The autofluorescence signals highlighted by the orange colour (Fig. S1e phasor pellet gradient bar) corresponds to a discrete, roughly spherical body (Figs. S1b and c), which we hypothesise to be the accumulation body. A single accumulation body is a hallmark feature of coccoid (<u>Freudenthal, 1962</u>) and dividing Symbiodiniaceae cells (<u>Camaya, 2020</u>) wherein only one of the mitotically dividing cells retains the accumulation body (<u>Camaya, 2020</u>; <u>Figueroa *et al*., 2021</u>). We found that most cells showed the presence of only one discrete accumulation body (orange in pseudo-coloured FLIM images), whereas in dividing cells there were two or more accumulation bodies. The accumulation body occasionally appeared in dividing cells to be dispersed throughout the cytoplasm, especially at 30 min post dawn (6:30 h) (Figs. 6k and l). The chemical composition of this body is unknown but it is reported to be adjacent to the nucleus and pyrenoid (<u>Camaya, 2020</u>; <u>Figueroa *et al*., 2021</u>; <u>Gornik *et al*., 2022</u>). This is consistent with the positioning of the accumulation body in our *B. minutum* cells (Fig. S8). A complete physio-chemical characterisation of the accumulation body in *B. minutum* is needed.

The pyrenoid, a site for carbon fixation that appears as a dense structure surrounded by a starch sheath, is reported to dissolve at night due to attenuation of CO_2_ fixation in the dark (<u>Griffiths, 1970</u>; <u>Atkinson *et al*., 2020</u>), and autofluorescence from this organelle is rarely reported unless excited by a 633 nm laser (<u>Uniacke *et al*., 2011</u>). We found that pyrenoid in *B. minutum* shows autofluorescence upon 488 excitation, however, signal intensity was lower compared to accumulation body.

The linear combination of multiexponential autofluorescence in *B. minutum* on phasor plots (Fig. 2f, orange pink coloured palette gradient bar) suggests a cumulative contribution from a several fluorescent chemicals possibly involved in energy quenching (<u>Subashchandrabose *et al*., 2014</u>). In other phototrophs, numerous autofluorescent pigments are involved in electron transport (as acceptors and donors) and are likely to quench fluorescence (<u>Broess *et al*., 2009</u>).

Various sample preparation approaches such as exposure of cells to sodium borohydride (<u>Castillo-Medina *et al*., 2011</u>), high light (<u>Webster *et al*., 2001</u>; <u>Maire *et al*., 2021</u>), combined hydrogen peroxide and light (<u>Yakubovskaya *et al*., 2019</u>), copper sulfate and ethanol (<u>Cohen *et al.*, 2021</u>), fluorescence lifetime gating (<u>Kodama, 2016</u>), TrueVIEW quenching kit (<u>Astafyeva *et al*., 2022b</u>) and post image processing using deep learning tools (<u>Jiang *et al*.,</u> <u>2022</u>) are used for reduction of background autofluorescence. <u>Maire *et al*. (2021)</u> quenched *B. minutum* cells by prolonged high light exposure (2 h, 400 µmole m^-2^ s^-1^), then FISH labelled and visualised ICB inside *B. minutum* cells. Photobleaching is the most straightforward process to reduce autofluorescence. However, high light exposure could not be used in this study exploring the impact of light on bacterial associations with *B. minutum*, since the light dictates algal cell stage transition (motile to coccoid) and division.

### Distinction of intracellular bacteria based on DY-490 lifetime signature

The presence of ICB is masked in CLSM images as the autofluorescence from the *B. minutum* overlaps with bacterial signals (Fig. 2c). The pseudo-colouring based on the fluorophore (DY-490) and autofluorescence specific cursor assignment in the phasor plot permits the visualisation of intracellular bacteria (Figs. 3f and i) using FLIM. The FLIM approach in this study used CLSM which facilitates visualisation of thin optical sections (0.7 µm), which is about one tenth of the *B. minutum* cell diameter (Fig 2c). Therefore, observation of bacteria in a given optical section is indeed present in intracellular spaces of *B. minutum* and are not an artifact.

### Spatial location of bacteria in *B. minutum*

FLIM images in this study demonstrate the presence of rod-shaped bacteria burrowing (Fig. 3i) into the cell exterior of *B. minutum*. SEM images (Fig. 3j–m) show rod- and spiral-shaped bacteria attached to the cell surface. This might suggest the existence of a specific receptor on the *B. minutum* surface which is recognised by bacterial cells and may act as a docking site or site of invasion by bacterial symbionts. <u>Astafyeva *et al*. (2022b)</u> reported invasion of the microalga, *Micrasterias radians* by the bacterium *Dyadobacter* sp. in a co-cultivation experiment. A similar exogenous uptake or invasion by bacteria in live *B. minutum* cells needs to be further explored. The higher percentage of *B. minutum* cells harbouring SAB compared with ICB, in light and dark conditions, suggests receptor-mediated bacterial attachment is likely. <u>Tran *et al*. (2022)</u> report that bacterial colonisation of the cell surface of the diatom, *Thalassiosira rotula*, is mediated by fucose-rich glycoconjugates. The *B. minutum* cell surface displays several glycoconjugates, D-mannose, D-glucose, lactose, D-galactose, L-fucose, N-acetylneuraminic acid and glucuronic acid (<u>Tortorelli *et al*., 2022</u>), however, their affinity for bacterial attachment has not been examined.

Maximum accumulation of ICB per *B. minutum* cell during the dark phase (22:30 h in Fig. 4) as observed in this study is corroborated by <u>Maire *et al*. (2021)</u> who reported a peak in % host population (*Cladocopium proliferum,* formerly referred to as *C. goreaui*) with closely associated bacteria (both SAB and ICB) at 22:00 h. The number of *C. proliferum* cells containing bacteria was 58% higher in the study by <u>Maire *et al*. (2021)</u> compared to *B. minutum* in this study. The former study used FISH and fluorescence activated cell sorting for estimation of host cell percentage containing ICB and/or SAB. This study provides a relative estimate of ICB only which was feasible due to the precise mapping of bacterial signals (based on DY-490) inside *B. minutum* by FLIM analysis. <u>Maire *et al*. (2021)</u> hypothesised that the increased percentage of Symbiodiniaceae (*C. proliferum*, *B. minutum* and *Fugacium* sp.) containing ICB in the dark phase is may be linked to heterotrophic feeding. <u>Jeong *et al*. (2012)</u> demonstrated mixotrophic mode of growth in *Symbiodinium* sp. with active ingestion of bacterial prey. Although a high ICB signal was observed under dark conditions, ∼60% of *B. minutum* cells were devoid of endosymbionts in this study. This suggest that the heterotrophic feeding behaviour (<u>Jeong *et al*., 2012</u>) was either absent or present in a very small subset of *B. minutum* population. Weak correlation between ICB signals per *B. minutum* cell in the dark and % *B. minutum* containing ICB suggest that the increase in intracellular bacterial load does not lead to more *B. minutum* cells harbouring the endosymbiotic bacteria. These bacterial symbionts are likely to be internalised by an unknown mechanism in the dark and symbiosis is facultative in nature. We observed the presence of bacteria in dividing furrows of *B. minutum* cells at the initiation of the light phase. We hypothesised that the re-organisation of thecal plates during cell division (<u>Kwok *et al.*, 2023</u>) in the dark is likely to result in internalisation of bacteria which is followed by de novo synthesis of the cell wall. The bacterial cells attached to the cell surface near the dividing furrow may become endosymbionts which is supported by the finding of diverse intracellular bacteria in this study. The motivation for bacteria to explore the *B. minutum* cell surface, such as via chemical attraction towards a nutrient rich phycosphere (<u>Garrido,</u> <u>Amana Guedes *et al*., 2021</u>), needs to be investigated.

### Diversity of intracellular bacteria in *B. minutum*

Similar to the previous report by <u>Maire *et al*. (2021)</u>, we report diverse communities of ICB with no differences in the light and dark phase. Many members of genera such as *Sphingomonas* (<u>Maire *et al*., 2021</u>), *Massilia* (<u>Sweet *et al*., 2021</u>), *Marinobacter (*<u>Sonnenschein *et al*., 2012</u>), *Endozoicomonas* (<u>Nishijima *et al*., 2013</u>) and *Methylobacterium* (<u>Green & Ardley, 2018</u>; <u>Maire *et al*., 2021</u>) that are found in the *B. minutum* microbiome are motile and/or rod-shaped. We hypothesise that the nutrient rich phycosphere (<u>Garrido,</u> <u>Amana Guedes *et al*., 2021</u>) of *B. minutum* can attract these motile bacteria which may incidentally result in a mutualistic association. *Marinobacter* sp. is one of the core microbiome members of *Breviolum* (<u>Lawson *et al*., 2018</u>) and a popular model system to study chemoattraction (<u>Garrido, Amana Guedes *et al*., 2021</u>). The motile and chemotactic behaviour of bacteria in the presence of *B. minutum* is, however, underexplored. The community composition of intracellular bacteria in *B. minutum* (SCF127-01, isolated from *Exapatisia diaphana* sourced from the Great Barrier Reef) in this study is strikingly different from that of *B. minutum* (MMSF01) examined in <u>Maire *et al*. (2021)</u>. Differences in the core microbiome of various Symbiodiniaceae genera are known (<u>Lawson *et al*., 2018</u>), however, variation in microbial genera at the *B. minutum* strain level have not been studied. These differences could arise from differences in the origin of *B. minutum*, i.e., host genotype that originally harboured the alga, and long-term culturing conditions impacting the microbial association.

Overall, the diversity of ICB, high ICB abundance in the dark independent % increase in *B. minutum* population with ICB, and the presence of a large portion of the *B. minutum* population lacking ICB suggest the *B. minutum*-bacteria symbiosis is non obligatory. Similarity in bacterial microbiome across different member of Symbiodiniaceae taxa is reported by Maire et al. (2022). The high bacterial diversity observed in this study suggest that a bacterial genus specific endosymbiosis as observed for *Micrasterias radians*-*Dyadobacter* sp. (<u>Astafyeva *et al*., 2022b</u>) is yet to be proven for *B. minutum*. Deciphering a specific symbiotic partner in *B. minutum* is essential for future studies (e.g., tracking of labelled bacteria) to advance our knowledge about processes involved in endosymbiont acquisition by this alga. There is a strong possibility that diverse bacteria may explore Symbiodiniaceae phycospheres as nutrient hotspot as observed in other microalgal (<u>Raina *et* *al.*, 2022</u>) and picophytoplankton species (<u>Raina *et al*., 2023</u>) which upon surface attachment or internalisation by an unknow mechanism results in epi and endosymbiosis. The consistent reporting of the presence of intracellular bacteria in this study and Maire et al. (2021) further raises questions about the metabolic exchange and reproduction of bacterial endosymbionts in *B. minutum*. We outline a novel FLIM based approach which enables future studies focused on entry and shading of bacterial symbiont, and spatial mapping of intracellular bacteria in *B. minutum*.

## Supporting information

Supplementary_sheet_S1

Supplementary_sheet_S2

## Acknowledgements

Authors acknowledge Gordon and Betty Moore foundation for provision of research funds (project no: 9351) to LLB, MvO, EH and DRB, and Dr(s). Jieqiong Lou, Gabriela Segal, and Allison van de Meene for training on FLIM, CLSM, and SEM (Biological Optical Microscopy Platform, University of Melbourne), respectively. MvO acknowledges Australian Research Council Laureate Fellowship FL180100036.

## Authors contribution

PD, MvO and LLB conceptualized the idea; PD performed experiment, analysed data and wrote the manuscript; SJTMC performed metabarcoding; DDB provided scientific inputs; EH provided inputs on FLIM data analysis; DDB, EH, MvO, LLB reviewed manuscript and obtained funding.

## Data availability

The raw metabarcoding reads and microscopy images generated and/or analysed during the current study are available in the Figshare repository, 10.6084/m9.figshare.24647403.

## References

Anthony CJ, Lock CC, Bentlage B. 2022. High-throughput physiological profiling of endosymbiotic dinoflagellates (Symbiodiniaceae) using flow cytometry. BioRxiv: 2022–2012.

Astafyeva Y, Gurschke M, Qi M, Bergmann L, Indenbirken D, de Grahl I, Katzowitsch E, Reumann S, Hanelt D, Alawi M. 2022a. Microalgae and bacteria interaction— Evidence for division of diligence in the alga microbiota. Microbiology Spectrum 10(4): e00633–00622.

Astafyeva Y, Gurschke M, Streit WR, Krohn I. 2022b. Interplay between the microalgae Micrasterias radians and its symbiont *Dyadobacter* sp. HH091. Frontiers in Microbiology 13: 4013.

Atkinson N, Mao Y, Chan KX, McCormick AJ. 2020. Condensation of Rubisco into a proto-pyrenoid in higher plant chloroplasts. Nature Communications 11(1): 1–9.

Bokulich NA, Kaehler BD, Rideout JR, Dillon M, Bolyen E, Knight R, Huttley GA, Gregory Caporaso J. 2018. Optimizing taxonomic classification of marker-gene amplicon sequences with QIIME 2’s q2-feature-classifier plugin. Microbiome 6(1): 1–17.

Bolyen E, Rideout JR, Dillon MR, Bokulich NA, Abnet CC, Al-Ghalith GA, Alexander H, Alm EJ, Arumugam M, Asnicar F. 2019. Reproducible, interactive, scalable and extensible microbiome data science using QIIME 2. Nature Biotechnology 37(8): 852–857.

Broess K, Borst JW, van Amerongen H. 2009. Applying two-photon excitation fluorescence lifetime imaging microscopy to study photosynthesis in plant leaves. Photosynthesis Research 100(2): 89–96.

Callahan BJ, McMurdie PJ, Rosen MJ, Han AW, Johnson AJA, Holmes SP. 2016. DADA2: High-resolution sample inference from Illumina amplicon data. Nature Methods 13(7): 581–583.

Camaya AP. 2020. Stages of the symbiotic zooxanthellae–host cell division and the dynamic role of coral nucleus in the partitioning process: A novel observation elucidated by electron microscopy. Coral Reefs 39(4): 929–938.

Castillo-Medina RE, Arzápalo-Castañeda G, Villanueva MA. 2011. Making walled-, highly autofluorescent dinoflagellate algae cells accessible and amenable for immunofluorescence and application of fluorescent probes. Limnology and Oceanography: Methods 9(10): 460–465.

Cohen AB, Novkov-Bloom A, Wesselborg C, Yagudaeva M, Aranguiz E, Taylor GT. 2021. Applying fluorescence in situ hybridization to aquatic systems with cyanobacteria blooms: autofluorescence suppression and high-throughput image analysis. Limnology and Oceanography: Methods 19(7): 457–475.

Davis NM, Proctor DM, Holmes SP, Relman DA, Callahan BJ. 2018. Simple statistical identification and removal of contaminant sequences in marker-gene and metagenomics data. Microbiome 6(1): 1–14.

Deore P, Wanigasuriya I, Ching SJTM, Brumley DR, van Oppen MJ, Blackall LL, Hinde E. 2022. Fluorescence lifetime imaging microscopy (FLIM): a non-traditional approach to study host-microbial symbioses. Microbiology Australia 43(1): 22–27.

Digman MA, Caiolfa VR, Zamai M, Gratton E. 2008. The phasor approach to fluorescence lifetime imaging analysis. Biophysical Journal 94(2): L14–16.

Figueroa R, Howe-Kerr L, Correa A. 2021. Direct evidence of sex and a hypothesis about meiosis in Symbiodiniaceae. Scientific Reports 11(1): 1–17.

Freudenthal HD. 1962. *Symbiodinium* gen. nov. and *Symbiodinium microadriaticum* sp. nov., a zooxanthella: taxonomy, life cycle, and morphology. The Journal of Protozoology 9(1): 45–52.

Frommlet JC, Wangpraseurt D, Sousa ML, Guimarães B, Medeiros da Silva M, Kühl M, Serôdio J. 2018. *Symbiodinium*-induced formation of microbialites: mechanistic insights from in vitro experiments and the prospect of its occurrence in nature. Frontiers in Microbiology 9: 998.

Gao C, Fernandez VI, Lee KS, Fenizia S, Pohnert G, Seymour JR, Raina J-B, Stocker R. 2020. Single-cell bacterial transcription measurements reveal the importance of dimethylsulfoniopropionate (DMSP) hotspots in ocean sulfur cycling. Nature Communications 11(1): 1–11.

Gardner SG, Leggat W, Ainsworth TD. 2023. The microbiome of the endosymbiotic Symbiodiniaceae in corals exposed to thermal stress. Hydrobiologia.

Garrido AG, Machado LF, Zilberberg C, Leite DCdA. 2021. Insights into ‘Symbiodiniaceae phycosphere’in a coral holobiont. Symbiosis 83(1): 25–39.

Gornik SG, Maegele I, Hambleton EA, Voss PA, Waller RF, Guse A. 2022. Nuclear transformation of a dinoflagellate symbiont of corals. Frontiers in Marine Science(9): 1035413.

Green PN, Ardley JK. 2018. Review of the genus *Methylobacterium* and closely related organisms: a proposal that some *Methylobacterium* species be reclassified into a new genus, Methylorubrum gen. nov. International Journal of Systematic and Evolutionary Microbiology 68(9): 2727–2748.

Griffiths DJ. 1970. The pyrenoid. The Botanical Review 36(1): 29–58.

Hartman LM. 2020. Manipulation of prokaryotic communities in the coral model organism, Exaiptasia diaphana. Ph. D. thesis, Swinburne University of Technology, Hawthorn.

Hinde E, Digman MA, Welch C, Hahn KM, Gratton E. 2012. Biosensor Förster resonance energy transfer detection by the phasor approach to fluorescence lifetime imaging microscopy. Microsc Res Tech 75(3): 271–281.

Jeong HJ, Yoo YD, Kang NS, Lim AS, Seong KA, Lee SY, Lee MJ, Lee KH, Kim HS, Shin W. 2012. Heterotrophic feeding as a newly identified survival strategy of the dinoflagellate *Symbiodinium*. Proceedings of the National Academy of Sciences 109(31): 12604–12609.

Jiang J, Zhang H, Orf GS, Lu Y, Xu W, Harrington LB, Liu H, Lo CS, Blankenship RE. 2014. Evidence of functional trimeric chlorophyll *a*/*c*_2_-peridinin proteins in the dinoflagellate *Symbiodinium*. Biochimica et Biophysica Acta (BBA)-Bioenergetics 1837(11): 1904–1912.

Jiang X, Pees T, Reinhold-Hurek B. 2022. Deep-learning-based removal of autofluorescence and fluorescence quantification in plant-colonizing bacteria inlZvivo. New Phytologist 235(6): 2481–2495.

Kato H, Tokutsu R, Kubota-Kawai H, Burton-Smith RN, Kim E, Minagawa J. 2020. Characterization of a Giant PSI Supercomplex in the Symbiotic Dinoflagellate Symbiodiniaceae. Plant Physiology 183(4): 1725–1734.

Kiene RP, Linn LJ, Bruton JA. 2000. New and important roles for DMSP in marine microbial communities. Journal of Sea Research 43(34): 209–224.

Kodama Y. 2016. Time gating of chloroplast autofluorescence allows clearer fluorescence imaging in planta. PloS One 11(3): e0152484.

Kwok ACM, Chan WS, Wong JTY. 2023. Dinoflagellate amphiesmal dynamics: cell wall deposition with ecdysis and cellular growth. Marine Drugs 21(2): 70.

Lawson CA, Raina JB, Kahlke T, Seymour JR, Suggett DJ. 2018. Defining the core microbiome of the symbiotic dinoflagellate, *Symbiodinium*. Environmental Microbiology Reports 10(1): 7–11.

Maire J, Girvan S, Barkla S, Perez-Gonzalez A, Suggett D, Blackall L, van Oppen M. 2021. Intracellular bacteria are common and taxonomically diverse in cultured and in hospite algal endosymbionts of coral reefs. The ISME Journal: 1–15.

Maire J, Tandon K, Collingro A, van de Meene A, Damjanovic K, Gotze CR, Stephenson S, Philip GK, Horn M, Cantin NE. 2022. *Endozoicomonas*-*chlamydiae* interactions in cell-associated microbial aggregates of the coral *Pocillopora acuta*. BioRxiv.

Martin M. 2011. Cutadapt removes adapter sequences from high-throughput sequencing reads. *EMBnet*. Journal 17(17): 10–12.

Matthews JL, Raina JB, Kahlke T, Seymour JR, van Oppen MJ, Suggett DJ. 2020. Symbiodiniaceae-bacteria interactions: rethinking metabolite exchange in reef-building corals as multi-partner metabolic networks. Environmental Microbiology 22(5): 1675–1687.

Motone K, Takagi T, Aburaya S, Miura N, Aoki W, Ueda M. 2020. A zeaxanthin-producing bacterium isolated from the algal phycosphere protects coral endosymbionts from environmental stress. mBio 11(1): e01019–01019.

Nishijima M, Adachi K, Katsuta A, Shizuri Y, Yamasato K. 2013. *Endozoicomonas numazuensis* sp. nov., a gammaproteobacterium isolated from marine sponges, and emended description of the genus *Endozoicomonas Kurahashi* and Yokota 2007. International Journal of Systematic and Evolutionary Microbiology 63(2): 709–714.

Nitschke MR, Fidalgo C, Simões J, Brandão C, Alves A, Serôdio J, Frommlet JC. 2020. Symbiolite formation: a powerful in vitro model to untangle the role of bacterial communities in the photosynthesis-induced formation of microbialites. The ISME journal 14(6): 1533–1546.

Periasamy A, Clegg R. 2009. FLIM microscopy in biology and medicine. USA: CRC Press.

Rädecker N, Pogoreutz C, Gegner HM, Cárdenas A, Perna G, Geißler L, Roth F, Bougoure J, Guagliardo P, Struck U, et al. 2022. Heat stress reduces the contribution of diazotrophs to coral holobiont nitrogen cycling. ISMEJ 16(4): 1110–1118.

Raina J-B, Giardina M, Brumley DR, Clode PL, Pernice M, Guagliardo P, Bougoure J, Mendis H, Smriga S, Sonnenschein EC, et al. 2023. Chemotaxis increases metabolic exchanges between marine picophytoplankton and heterotrophic bacteria. Nature Microbiology 8(3): 510–521.

Raina J-B, Lambert BS, Parks DH, Rinke C, Siboni N, Bramucci A, Ostrowski M, Signal B, Lutz A, Mendis H, et al. 2022. Chemotaxis shapes the microscale organization of the ocean’s microbiome. Nature 605(7908): 132–138.

Raina JB, Dinsdale EA, Willis BL, Bourne DG. 2010. Do the organic sulfur compounds DMSP and DMS drive coral microbial associations? Trends in Microbiology 18(3): 101–108.

Ranjit S, Malacrida L, Jameson DM, Gratton E. 2018. Fit-free analysis of fluorescence lifetime imaging data using the phasor approach. Nature Protocol 13(9): 1979–2004.

Sonnenschein EC, Syit DA, Grossart H-P, Ullrich MS. 2012. Chemotaxis of *Marinobacter adhaerens* and its impact on attachment to the diatom *Thalassiosira weissflogii*. Applied and Environmental Microbiology 78(19): 6900–6907.

Subashchandrabose SR, Krishnan K, Gratton E, Megharaj M, Naidu R. 2014. Potential of fluorescence imaging techniques to monitor mutagenic PAH uptake by microalga. Environmental Science & Technology 48(16): 9152–9160.

Sweet M, Villela H, Keller-Costa T, Costa R, Romano S, Bourne DG, Cárdenas A, Huggett MJ, Kerwin AH, Kuek F, et al. 2021. Insights into the Cultured Bacterial Fraction of Corals. mSystems 6(3): e01249–01220.

Tang YZ, Dobbs FC. 2007. Green autofluorescence in dinoflagellates, diatoms, and other microalgae and its implications for vital staining and morphological studies. Applied and Environmental Microbiology 73(7): 2306–2313.

Tortorelli G, Rautengarten C, Bacic A, Segal G, Ebert B, Davy SK, van Oppen MJ, McFadden GI. 2022. Cell surface carbohydrates of symbiotic dinoflagellates and their role in the establishment of cnidarian–dinoflagellate symbiosis. The ISME journal 16(1): 190– 199.

Tran QD, Neu TR, Sultana S, Giebel HA, Simon M, Billerbeck S. 2022. Distinct glycoconjugate cell surface structures make the pelagic diatom *Thalassiosira rotula* an attractive habitat for bacteria. Journal of Phycology 59(2): 309–322.

Uniacke J, Colón-Ramos D, Zerges W 2011. FISH and immunofluorescence staining in *Chlamydomonas*. In: Gerst JE ed. RNA Detection and Visualization. Totowa, NJ: Humana Press, 15–29.

Wakefield TS, Farmer MA, Kempf SC. 2000. Revised description of the fine structure of in situ" zooxanthellae" genus *Symbiodinium*. The Biological Bulletin 199(1): 76–84.

Wang LH, Liu YH, Ju YM, Hsiao YY, Fang LS, Chen CS. 2008. Cell cycle propagation is driven by light–dark stimulation in a cultured symbiotic dinoflagellate isolated from corals. Coral Reefs 27(4): 823–835.

Webster NS, Wilson KJ, Blackall LL, Hill RT. 2001. Phylogenetic diversity of bacteria associated with the marine sponge *Rhopaloeides odorabile*. Applied and Environmental Microbiology 67(1): 434–444.

Yakubovskaya E, Zaliznyak T, Martínez Martínez J, Taylor GT. 2019. Tear down the fluorescent curtain: a new fluorescence suppression method for Raman microspectroscopic analyses. Scientific Reports 9(1): 1–9.

Yonge C. 1931. Studies on the Physiology of Corals IV. The Structure Distribution and Physiology of the Zooxanthellae. . Scientific Reports 1: 135–176.

