## Supplementary_sheet_S1 for "Cutting through host autofluorescence: fluorescence lifetime imaging microscopy for visualising intracellular bacteria in Symbiodiniaceae"

### Material and methods:

#### 16S rRNA gene metabarcoding

#### DNA extraction, amplification, and library preparation

DNA was extracted from biomass on filters using a method outlined by Wilson et al. (2002). Briefly, 0.375 mL of freshly prepared filter sterilised (0.22 µm) extraction buffer (Tris buffer pH-9, 100 mM EDTA, 100 mM NaCl; 1% SDS, 0.2 mg $mL^{-1}$ lysozyme) was added to a 1.5 mL microcentrifuge tube with the biomass-containing filters. Tubes were vortexed and incubated at 37°C for 30 min. Tubes containing only extraction buffer served as extraction blanks (n = 3) to ensure DNA preparation was free of contamination from chemicals. Proteinase K at a final concentration of 0.5 mg $mL^{-1}$ was added to digest nucleases and other proteins. Samples were subjected to mechanical cell lysis using ~100 mg sterile glass beads (425–600 µm diameter) followed by bead beating for 30 s at 30 Hz on a Qiagen tissue-lyser II. Samples were incubated at 65°C for 60 min. DNA was precipitated using 0.45 mM ice-cold potassium acetate followed by 30 min of incubation on ice. Samples were centrifuged twice at 15000 x *g* for 5 min and the supernatant was treated with RNAse (final concentration 0.13 mg $mL^{-1}$) at 37°C for 30 min. DNA was further purified and precipitated using 500 µL of ice-cold iso-propanol, samples were mixed gently and incubated at room temperature (RT) for 15 min. DNA was pelleted by centrifuging samples at 15000 x *g* for 15 min. The DNA pellets were washed twice with ice-cold ethanol and centrifuged at 15000 x *g* for 2 min. All samples were air dried at RT for ~40 min and DNA was resuspended in MilliQ water.

The ICB community composition was characterised in bulk Symbiodiniaceae cells using 16S rRNA gene metabarcoding, in which triplicates of each sample were amplified using 16S rRNA gene primers spanning hypervariable V5 and V6 regions (784f – 5' GTGACCTATGAACTCAGGAGTCAGGATTAGATACCCTGGTA and 1061r – 5' CTGAGACTTGCACATCGCAGCCRRCACGAGCTGACGAC 3'; underlined nucleotides indicate Illumina adaptors). The PCR mix contained 1X MyTaq mix, 0.2 µm of 784f and 1061r, variable concentrations of DNA template (42%, 62%, 96% or 100% v/v) and nuclease-free water at a volume to make up the total reaction volume to 50 µL. Variable template DNA concentrations were used in overhang PCRs because occasionally the DNA yield was very low from sodium hypochlorite-washed biomass (see supplementary sheet S1). A no template DNA served as a negative PCR control (n = 5) to detect any potential contaminants arising from the PCR kit. The amplification settings were 95°C for 3 min for initial denaturation followed by 18 cycles of denaturation at 95°C for 15 s, annealing at 55°C for 30 s, extension at 95°C for 30 s, and a final extension at 72°C for 7 min. Three set of PCRs using above conditions were performed which yielded triplicates of each sample that was treated as technical replicates with distinct barcode. A separate PCR of 35 cycles using identical denaturation, annealing and extension conditions were performed to check for the expected size amplicon (277 bp) by gel electrophoresis. For library preparation, bead clean-up (0.8 beads-to-PCR product ratio) was performed on pooled samples which were prepared by mixing of 5 µL of PCR product from technical triplicates of each sample followed by washing of beads twice with 150 µL of 70% ethanol. The adsorbed DNA from the beads were released by adding 40 µL of nuclease-free $H_2O$. The clean-up beads and indexing primers were obtained from Water and Eliza Hall Institute (WEHI), Melbourne, Australia.

Indexing PCR was performed using 10 µL cleaned DNA suspended in nuclease-free $H_2O$ as a template, 0.25 µM of forward and reverse indexing primers, and 1X of MyTaq PCR mix. The amplification conditions were an initial denaturation cycle at 95°C for 3 min, 24 cycles of denaturation at 95°C for 15 s, annealing at 60°C for 30 s, extension at 72°C for 30 s and final extension at 72°C for 7 min. An aliquot of 5 µL from each technical replicate, extraction blank and no DNA template control was pooled which was mixed with 30 µL of NGS beads and incubated at RT for 5 min. These beads were washed twice with 180 µL of 80% ethanol and air dried. DNA was released from the beads using nuclease-free water and samples were sequenced on a single illumina MiSeq run using v3 (2 x 300 bp) at WEHI.

**Statistical analysis of ASVs and visualisation in R studio**

Samples with highest number of reads in each replicate samples were considered for ASV analysis and visualisation because of variation in template DNA concentration (42, 62, 96 or 100%) used in PCR. No particular DNA concentration yielded a consistent PCR amplification across all replicates. We believe that pooling of sub-replicate samples that were amplified using variable DNA concentrations may have introduced bias, for example, over-representation of certain bacterial taxa. Sequencing PCR products from only one set of DNA concentration on occasion would have resulted in loss of replicates of a sample for some time-points. Therefore, all samples (4 DNA concentration x 8 timepoints x 5 replicate for each timepoint, n = 160) were sequenced. All samples were post processed in QIIME 2.2022-8. For each timepoints, samples yielding highest number of ASVs regardless of DNA concentration were used for downstream analysis in R. We have provided visualisation of relative abundances of bacterial ASV and beta-diversity analysis across 160 samples in supplementary sheet S2.

**Results**:

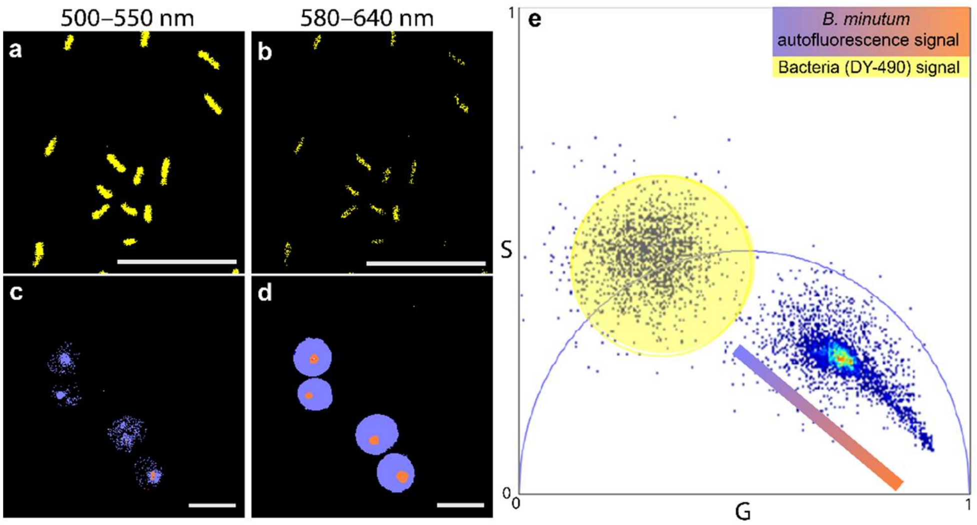


Fig. S1. Fluorescence lifetime imaging microscopy images of a, b) pure culture of *Marinobacter adhaerens* labelled with EUB338-DY-490, and c, d) *Breviolum minutum* autofluorescence. ex 488 nm, em 500–550 nm and em 580–640 nm as indicated above the panels. e) A phasor plot indicating signal counts of DY-490 in large yellow coloured cursor (exhibiting single exponential lifetime of ~2.9 ns) placed on phasor coordinates s, g (0.571,

0.236) falling on the line of universal circle The linear combinations of *B. minutum* specific fluorescence multi-exponential lifetimes are presented as blue to orange coloured pellet. The colours (yellow, blue and orange coloured palette for gradient bar) in phasor plot used for pseudo-colouring of images show in a–d.

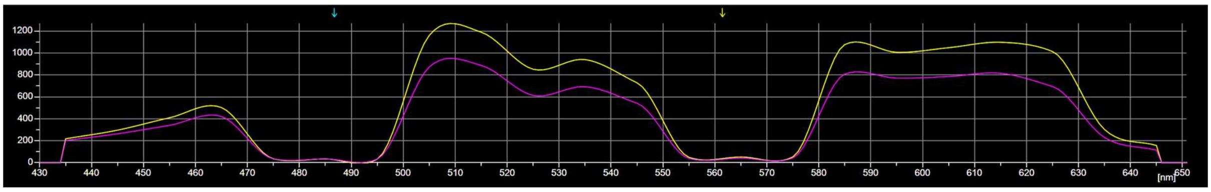


Fig. S2. Autofluorescence spectra of *B. minutum* cells fixed in 4% (v/v) paraformaldehyde. Excitation lasers were 409, 488 (top cyan arrow) and 561 (top green arrow) nm, and yellow and pink lines indicates signals from two *B. minutum* cells.

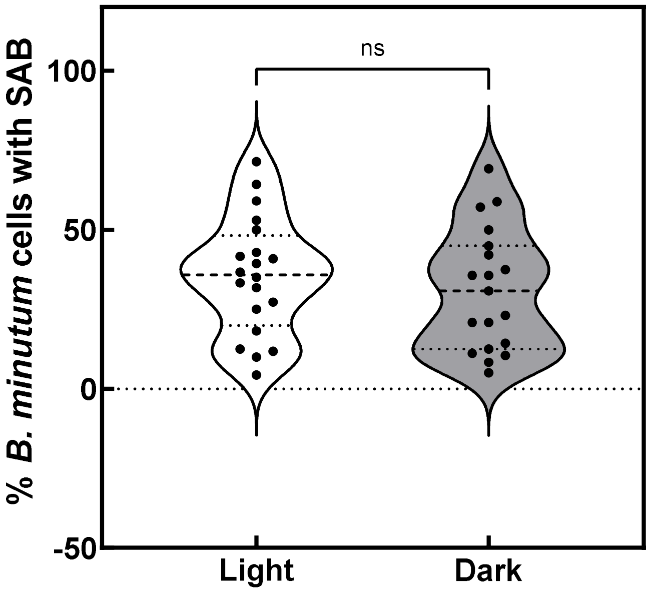


Fig. S3. Proportion of *B. minutum* cells with surface attached bacteria (SAB) in light and dark phase. Differences are non-significant (ns) at the statistical level of significance 0.05. Welch's t-test t = 0.736, df 36.75.

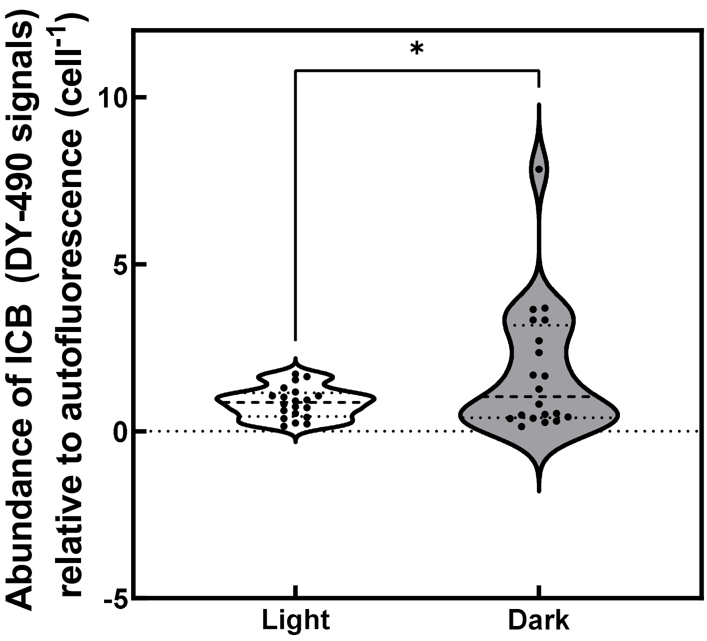


Fig. S4. Relative abundance of intracellular bacteria (ICB) in *B. minutum* cell based on feature counts of DY-490 in phasor plot under light and dark phase. p value * is 0.0453, Welch's t-test (t=2.127, df=21.27) where statistical level of significance was 0.05.

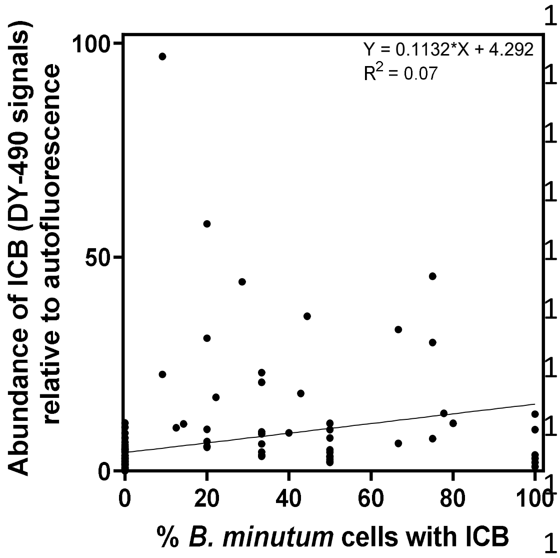


Fig. S5. Correlation between abundance of ICB normalised to autofluorescence (per cell) and percentage of *B. minutum* population containing ICB. Data point indicate mean values in independent biological replicates (n = 5).

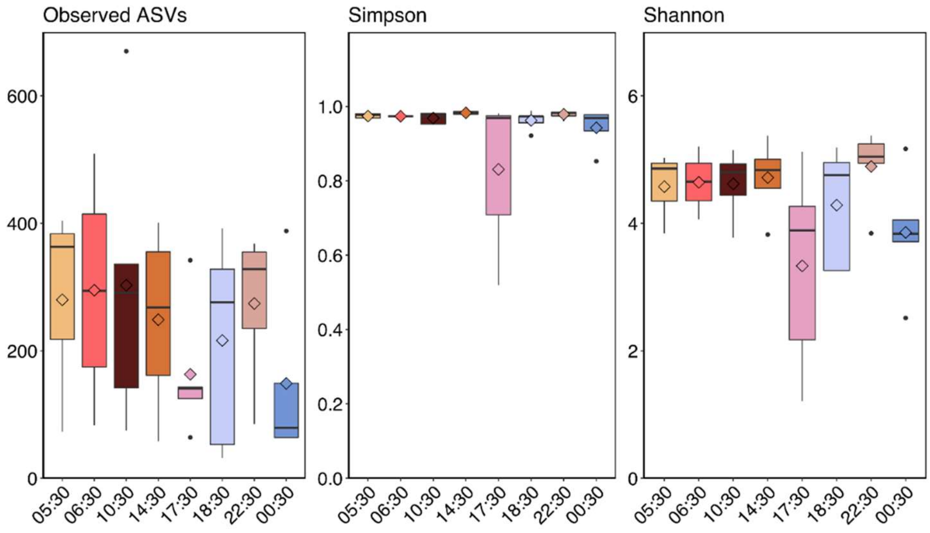


Fig. S6. Alpha diversity of intracellular bacteria in *B. minutum* based on Observed ASVs, Simpson and Shannon indices under light (6:30–17:30 h) and dark (18:30–05:30 h) conditions.

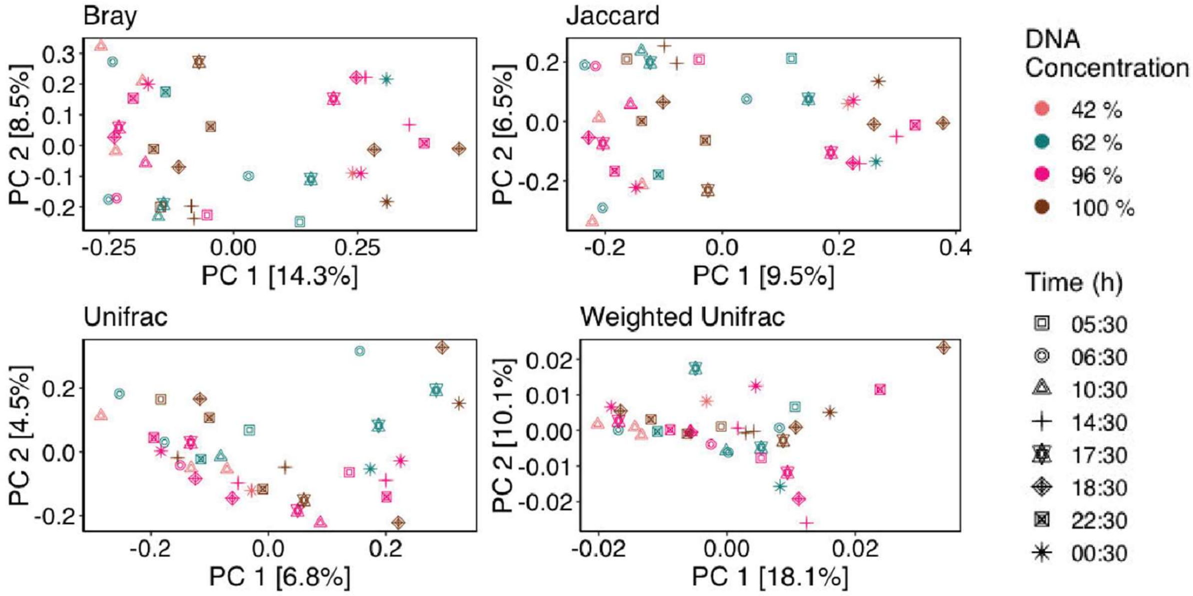


Fig. S7. Differences in intracellular microbiome of *B. minutum* based on dissimilarity matrices (beta diversity) in light (6:30–17:30 h) and dark (18:30–05:30 h) phase (black symbol legends), and in samples that were amplified using variable amounts of template DNA (coloured circle legends). Data is analysed from 40 samples (8 timepoints x 5 replicates)

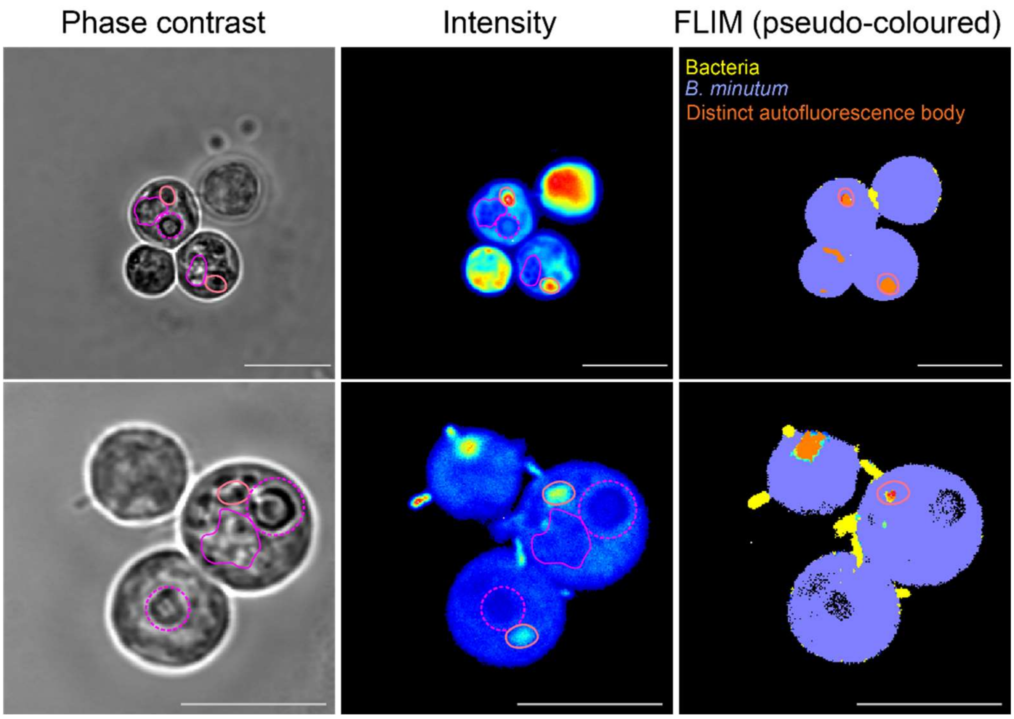


Fig. S8. Positioning of *B. minutum* organelles within the algal cell. Nuclei (pink irregular shape), accumulation body (orange circle in phase and intensity images, solid orange body in FLIM images) and pyrenoid (dashed pink circles). Scale bar = 10 µm.
