## Supplementary_sheet_S2 for "Cutting through host autofluorescence: fluorescence lifetime imaging microscopy for visualising intracellular bacteria in Symbiodiniaceae"

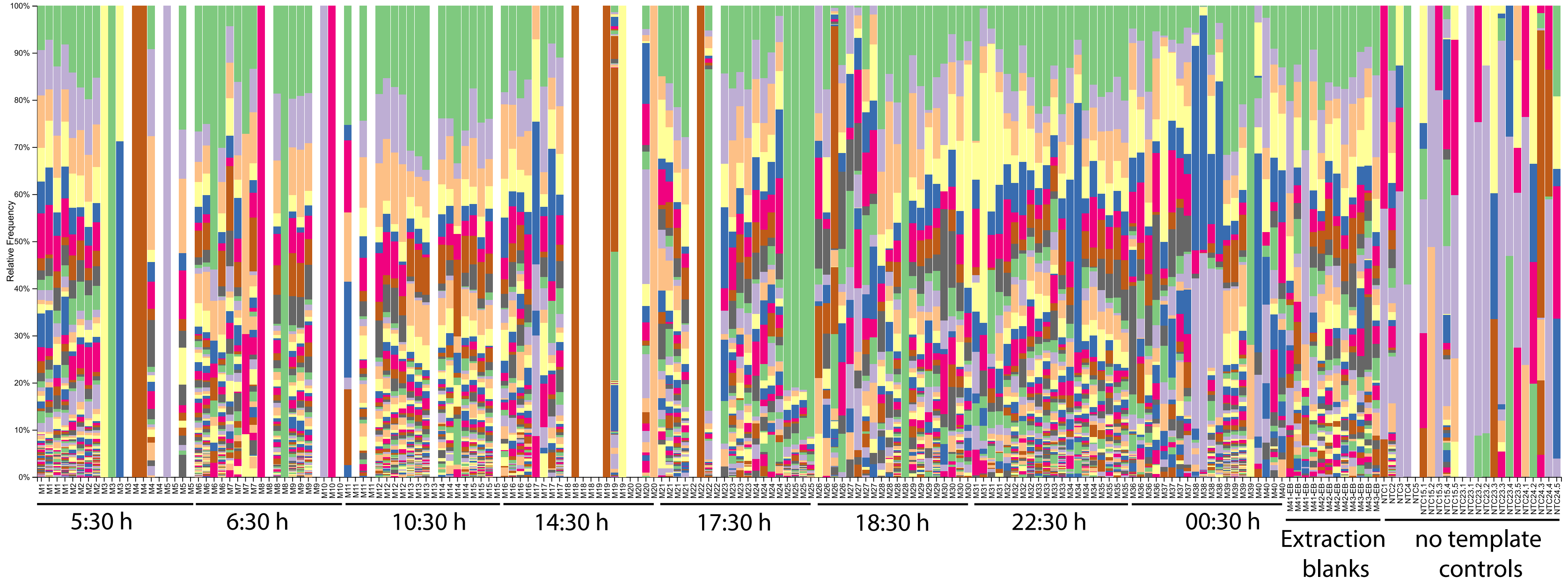
100%
90%
80%
70%
60%
50%
40%
30%
20%
10%
0%
Relative Frequency
5:30 h
6:30 h
10:30 h
14:30 h
17:30 h
18:30 h
22:30 h
00:30 h
Extraction blanks
no template controls

d__Bacteria;p__Actinobacteriota;c__Actinobacteria;o__Micrococcales
d__Bacteria;p__Bacteroidota;c__Bacteroidia;o__Flavobacteriales
d__Bacteria;p__Proteobacteria;c__Alphaproteobacteria;o__Rhodobacterales
d__Bacteria;p__Proteobacteria;c__Alphaproteobacteria;o__Rhizobiales
d__Bacteria;p__Proteobacteria;c__Gammaproteobacteria;o__Burkholderiales
d__Bacteria;p__Proteobacteria;c__Alphaproteobacteria;o__Sphingomonadales
d__Bacteria;p__Bacteroidota;c__Bacteroidia;o__Cytophagales
d__Bacteria;p__Proteobacteria;c__Gammaproteobacteria;o__Oceanospirillales
d__Bacteria;p__Actinobacteriota;c__Actinobacteria;o__Propionibacteriales
d__Bacteria;p__Proteobacteria;c__Gammaproteobacteria;o__Pseudomonadales
d__Bacteria;p__Proteobacteria;c__Gammaproteobacteria;o__Alteromonadales
d__Bacteria;p__Proteobacteria;c__Alphaproteobacteria;o__Rickettsiales
d__Bacteria;p__Proteobacteria;c__Alphaproteobacteria;o__Caulobacterales
d__Bacteria;p__Firmicutes;c__Bacilli;o__Staphylococcales
d__Bacteria;p__Bacteroidota;c__Bacteroidia;o__Bacteroidales
d__Bacteria;p__Firmicutes;c__Bacilli;o__Bacillales
d__Bacteria;p__Bacteroidota;c__Bacteroidia;o__Chitinophagales
d__Bacteria;p__Actinobacteriota;c__Actinobacteria;o__Corynebacteriales
d__Bacteria;p__Firmicutes;c__Bacilli;o__Lactobacillales
d__Bacteria;p__Proteobacteria;c__Gammaproteobacteria;o__Cellvibrionales
d__Bacteria;p__Planctomycetota;c__Phycisphaerae;o__Phycisphaerales
d__Bacteria;p__Proteobacteria;c__Gammaproteobacteria;o__Xanthomonadales
d__Bacteria;p__Firmicutes;c__Clostridia;o__Lachnospirales
d__Bacteria;p__Proteobacteria;c__Gammaproteobacteria;__
d__Bacteria;p__Firmicutes;c__Clostridia;o__Oscillospirales
d__Bacteria;p__Proteobacteria;c__Gammaproteobacteria;o__Steroidobacterales
d__Bacteria;p__Proteobacteria;c__Gammaproteobacteria;o__Enterobacterales
d__Bacteria;p__Verrucomicrobiota;c__Chlamydiae;o__Chlamydiales
d__Bacteria;p__Cyanobacteria;c__Cyanobacteriia;o__Chloroplast
d__Bacteria;p__Proteobacteria;c__Alphaproteobacteria;o__Kiloniellales
d__Bacteria;p__Bacteroidota;c__Bacteroidia;o__Sphingobacteriales
d__Bacteria;p__Proteobacteria;c__Alphaproteobacteria;__
d__Bacteria;p__Proteobacteria;c__Alphaproteobacteria;o__Rhodospirillales
d__Bacteria;p__Proteobacteria;c__Alphaproteobacteria;o__Parvibaculales
d__Bacteria;p__Proteobacteria;c__Gammaproteobacteria;o__Vibrionales
d__Bacteria;p__Firmicutes;c__Clostridia;o__Peptostreptococcales-Tissierellales

d__Bacteria;p__Fusobacteriota;c__Fusobacteriia;o__Fusobacteriales
d__Bacteria;__;__;__
d__Bacteria;p__Proteobacteria;c__Gammaproteobacteria;o__KI89A_clade
d__Bacteria;p__Proteobacteria;c__Alphaproteobacteria;o__Thalassobaculales
d__Bacteria;p__NB1-j;c__NB1-j;o__NB1-j
d__Bacteria;p__Proteobacteria;c__Alphaproteobacteria;o__Micavibrionales
d__Bacteria;p__Proteobacteria;c__Gammaproteobacteria;o__Gammaproteobacteria_Incertae_Sedis
d__Bacteria;p__Proteobacteria;__;__
d__Bacteria;p__Proteobacteria;c__Gammaproteobacteria;o__Tenderiales
d__Bacteria;p__Proteobacteria;c__Gammaproteobacteria;o__Salinisphaerales
d__Bacteria;p__Bacteroidota;c__Rhodothermia;o__Balneolales
d__Bacteria;p__Proteobacteria;c__Alphaproteobacteria;o__uncultured
d__Bacteria;p__Firmicutes;c__Bacilli;o__Paenibacillales
d__Bacteria;p__Bdellovibrionota;c__Bdellovibrionia;o__Bacteriovoracales
d__Bacteria;p__Myxococcota;c__Polyangia;o__Nannocystales
d__Bacteria;p__Actinobacteriota;c__Acidimicrobiia;o__Microtrichales
d__Bacteria;p__Firmicutes;c__Clostridia;o__Clostridia_UCG-014
d__Bacteria;p__Myxococcota;c__Polyangia;o__Polyangiales
d__Bacteria;p__Actinobacteriota;c__Actinobacteria;o__Actinomycetales
d__Bacteria;p__Acidobacteriota;c__Acidobacteriae;o__Subgroup_2
d__Bacteria;p__Bdellovibrionota;c__Bdellovibrionia;o__Bdellovibrionales
d__Bacteria;p__Proteobacteria;c__Gammaproteobacteria;o__OM182_clade
d__Bacteria;p__Proteobacteria;c__Alphaproteobacteria;o__Acetobacterales
d__Bacteria;p__Proteobacteria;c__Gammaproteobacteria;o__Thiohalorhabdales
d__Bacteria;p__Acidobacteriota;c__Thermoanaerobaculia;o__Thermoanaerobaculales
d__Bacteria;p__Spirochaetota;c__Spirochaetia;o__Spirochaetales
d__Bacteria;p__Bdellovibrionota;c__Oligoflexia;o__0319-6G20
d__Bacteria;p__Planctomycetota;c__Planctomycetes;o__Pirellulales
d__Bacteria;p__Actinobacteriota;c__Actinobacteria;o__Frankiales
d__Bacteria;p__Proteobacteria;c__Gammaproteobacteria;o__Francisellales
d__Bacteria;p__Campilobacterota;c__Campylobacteria;o__Campylobacterales
d__Bacteria;p__Proteobacteria;c__Gammaproteobacteria;o__Orbales
d__Bacteria;p__Verrucomicrobiota;c__Verrucomicrobiae;o__Pedosphaerales
d__Bacteria;p__Proteobacteria;c__Gammaproteobacteria;o__CHAB-XI-27
d__Bacteria;p__Proteobacteria;c__Alphaproteobacteria;o__NRL2
d__Bacteria;p__Planctomycetota;c__OM190;o__OM190

[illegible]
d__Bacteria;p__Proteobacteria;c__Alphaproteobacteria;o__Dstr-E11
d__Bacteria;p__Proteobacteria;c__Gammaproteobacteria;o__JG36-TzT-191
d__Bacteria;p__Proteobacteria;c__Alphaproteobacteria;o__Kordiimonadales
d__Bacteria;p__Bdellovibrionota;c__Oligoflexia;o__Oligoflexales
d__Bacteria;p__Actinobacteriota;c__Actinobacteria;o__Pseudonocardiales
d__Bacteria;p__Dependentiae;c__Babeliae;o__Babeliales
d__Bacteria;p__Desulfobacterota;c__Desulfovibrionia;o__Desulfovibrionales
d__Bacteria;p__Proteobacteria;c__Gammaproteobacteria;o__Legionellales
d__Bacteria;p__Proteobacteria;c__Alphaproteobacteria;o__Sneathiellales
d__Bacteria;p__Myxococcota;c__Polyangia;o__Haliangiales
d__Bacteria;p__Planctomycetota;c__Planctomycetes;o__Planctomycetales
d__Bacteria;p__Proteobacteria;c__Gammaproteobacteria;o__Diplorickettsiales
d__Bacteria;p__Spirochaetota;c__Leptospirae;o__Leptospirales
d__Bacteria;p__Firmicutes;c__Clostridia;o__Eubacteriales
d__Bacteria;p__Proteobacteria;c__Alphaproteobacteria;o__Reyranellales
d__Bacteria;p__Proteobacteria;c__Gammaproteobacteria;o__Nitrosococcales
d__Bacteria;p__Bacteroidota;c__Bacteroidia;__
d__Bacteria;p__Actinobacteriota;c__Actinobacteria;o__Bifidobacteriales
d__Bacteria;p__Bacteroidota;c__Rhodothermia;o__Rhodothermales
d__Bacteria;p__SAR324_clade(Marine_group_B);c__SAR324_clade(Marine_group_B);o__SAR324_clade(Marine_group_B)
d__Bacteria;p__Planctomycetota;c__BD7-11;o__BD7-11
d__Bacteria;p__Acidobacteriota;c__Subgroup_22;o__Subgroup_22
d__Bacteria;p__Actinobacteriota;c__Thermoleophilia;o__Gaiellales
d__Bacteria;p__Firmicutes;c__Bacilli;o__Thermicanales
d__Bacteria;p__Bacteroidota;c__Kapabacteria;o__Kapabacteriales
d__Bacteria;p__Firmicutes;c__Clostridia;o__Clostridia_vadinBB60_group
d__Bacteria;p__Actinobacteriota;c__Coriobacteriia;o__Coriobacteriales
d__Bacteria;p__Proteobacteria;c__Gammaproteobacteria;o__Arenicellales
d__Bacteria;p__Proteobacteria;c__Gammaproteobacteria;o__pItb-vmat-80
d__Bacteria;p__Firmicutes;c__Bacilli;o__Brevibacillales
d__Bacteria;p__Acidobacteriota;c__Aminicenantia;o__Aminicenantales
d__Bacteria;p__Myxococcota;c__Myxococcia;o__Myxococcales
d__Bacteria;p__Proteobacteria;c__Alphaproteobacteria;o__Defluviicoccales
d__Bacteria;p__Proteobacteria;c__Gammaproteobacteria;o__Ga0077536
d__Bacteria;p__Proteobacteria;c__Gammaproteobacteria;o__MD2904-B13
[illegible]

d__Bacteria;p__Gemmatimonadota;c__BD2-11_terrestrial_group;o__BD2-11_terrestrial_group
d__Bacteria;p__Firmicutes;c__Bacilli;o__Erysipelotrichales
d__Bacteria;p__Proteobacteria;c__Gammaproteobacteria;o__HOC36
d__Bacteria;p__Acidobacteriota;c__Holophagae;o__Acanthopleuribacterales
d__Bacteria;p__Armatimonadota;c__Fimbriimonadia;o__Fimbriimonadales
d__Bacteria;p__Actinobacteriota;c__Actinobacteria;o__Micromonosporales
d__Bacteria;p__Firmicutes;c__Clostridia;o__Monoglobales
d__Bacteria;p__Myxococcota;c__Polyangia;o__mle1-27
d__Bacteria;p__Proteobacteria;c__Alphaproteobacteria;o__Puniceispirillales
d__Bacteria;p__Proteobacteria;c__Gammaproteobacteria;o__JTB23
d__Bacteria;p__Proteobacteria;c__Alphaproteobacteria;o__Elsterales
d__Bacteria;p__Desulfobacterota;c__Desulfuromonadia;o__PB19
d__Bacteria;p__Firmicutes;c__Clostridia;o__Christensenellales
d__Bacteria;p__Proteobacteria;c__Gammaproteobacteria;o__UBA10353_marine_group
d__Bacteria;p__Planctomycetota;c__Pla3_lineage;o__Pla3_lineage
d__Bacteria;p__Proteobacteria;c__Gammaproteobacteria;o__UBA4486
d__Bacteria;p__Proteobacteria;c__Alphaproteobacteria;o__Tistrellales
d__Bacteria;p__Proteobacteria;c__Alphaproteobacteria;o__Ferrovibrionales
d__Bacteria;p__Proteobacteria;c__Gammaproteobacteria;o__AT-s2-59
d__Bacteria;p__Desulfobacterota;c__Desulfuromonadia;o__Bradymonadales
d__Bacteria;p__Proteobacteria;c__Alphaproteobacteria;o__Micropepsales
d__Bacteria;p__Proteobacteria;c__Gammaproteobacteria;o__Pasteurellales
d__Bacteria;p__Proteobacteria;c__Gammaproteobacteria;o__eub62A3
d__Bacteria;p__Firmicutes;c__Clostridia;o__Clostridiales
d__Bacteria;p__Proteobacteria;c__Gammaproteobacteria;o__uncultured
d__Bacteria;p__Actinobacteriota;c__Acidimicrobiia;o__Actinomarinales
d__Bacteria;p__Acidobacteriota;c__Acidobacteriae;o__Bryobacterales
d__Bacteria;p__Proteobacteria;c__Alphaproteobacteria;o__Azospirillales
d__Bacteria;p__Proteobacteria;c__Alphaproteobacteria;o__Paracaedibacterales
d__Bacteria;p__Proteobacteria;c__Gammaproteobacteria;o__MBAE14
d__Bacteria;p__Proteobacteria;c__Gammaproteobacteria;o__SZB50
d__Bacteria;p__Proteobacteria;c__Gammaproteobacteria;o__Beggiatoales
d__Bacteria;p__Desulfobacterota;c__uncultured;o__uncultured
d__Bacteria;p__Acidobacteriota;c__Subgroup_26;o__Subgroup_26
d__Bacteria;p__Patescibacteria;c__Gracilibacteria;o__Absconditabacteriales_(SR1)
Unassigned;__;__;__

d__Bacteria;p__Firmicutes;c__Bacilli;__
d__Bacteria;p__Hydrogenedentes;c__Hydrogenedentia;o__Hydrogenedentiales
d__Bacteria;p__Calditrichota;c__Calditrichia;o__Calditrichales
d__Bacteria;p__Proteobacteria;c__Gammaproteobacteria;o__SAR86_clade
d__Bacteria;p__Proteobacteria;c__Gammaproteobacteria;o__Coxiellales
d__Bacteria;p__Actinobacteriota;c__Thermoleophilia;o__Solirubrobacterales
d__Bacteria;p__Patescibacteria;c__Gracilibacteria;o__Gracilibacteria
d__Bacteria;p__Acidobacteriota;c__Acidobacteriae;o__Acidobacteriales
d__Bacteria;p__Firmicutes;c__Thermoanaerobacteria;o__Thermoanaerobacterales
d__Bacteria;p__Firmicutes;c__Clostridia;o__MAT-CR-H4-C10
d__Bacteria;p__Elusimicrobiota;c__Elusimicrobia;o__Lineage_IV
d__Bacteria;p__Proteobacteria;c__Gammaproteobacteria;o__Granulosicoccales
d__Bacteria;p__Myxococcota;c__Polyangia;o__UASB-TL25
d__Bacteria;p__Proteobacteria;c__Gammaproteobacteria;o__Ectothiorhodospirales
d__Bacteria;p__Actinobacteriota;c__Actinobacteria;o__Streptomycetales
d__Bacteria;p__Desulfobacterota;c__Desulfuromonadia;o__Geobacterales
d__Bacteria;p__Proteobacteria;c__Gammaproteobacteria;o__Thiotrichales
d__Bacteria;p__Proteobacteria;c__Gammaproteobacteria;o__EC3
d__Bacteria;p__Firmicutes;c__Desulfotomaculia;o__Desulfotomaculales
d__Bacteria;p__Chloroflexi;c__Anaerolineae;o__Ardenticatenales
d__Bacteria;p__Proteobacteria;c__Gammaproteobacteria;o__Gammaproteobacteria
d__Bacteria;p__Proteobacteria;c__Gammaproteobacteria;o__CCM19a
d__Bacteria;p__Patescibacteria;c__Saccharimonadia;o__Saccharimonadales
d__Bacteria;p__Verrucomicrobiota;c__Lentisphaeria;o__Victivallales
d__Bacteria;p__MBNT15;c__MBNT15;o__MBNT15
d__Bacteria;p__Gemmatimonadota;c__Gemmatimonadetes;o__Gemmatimonadales
d__Bacteria;p__Myxococcota;c__Polyangia;o__Blfdi19
d__Bacteria;p__Planctomycetota;c__Phycisphaerae;o__CCM11a
d__Bacteria;p__Acidobacteriota;c__Acidobacteriae;__
d__Bacteria;p__Firmicutes;c__Clostridia;__
d__Bacteria;p__Proteobacteria;c__Alphaproteobacteria;o__Holosporales
d__Bacteria;p__Proteobacteria;c__Gammaproteobacteria;o__SS1-B-07-19
d__Bacteria;p__Proteobacteria;c__Gammaproteobacteria;o__Methylococcales
d__Bacteria;p__Firmicutes;c__Bacilli;o__Exiguobacterales
d__Bacteria;p__Myxococcota;c__bacteriap25;o__bacteriap25
d__Bacteria;p__Actinobacteriota;c__Actinobacteria;__

d__Bacteria;p__PAUC34f;c__PAUC34f;o__PAUC34f
d__Bacteria;p__Firmicutes;__;__
d__Bacteria;p__Proteobacteria;c__Gammaproteobacteria;o__EPR3968-O8a-Bc78
d__Bacteria;p__Actinobacteriota;c__Actinobacteria;o__Kineosporiales
d__Bacteria;p__Patescibacteria;c__Gracilibacteria;o__Candidatus_Peribacteria
d__Bacteria;p__Actinobacteriota;c__Acidimicrobiia;o__IMCC26256
d__Bacteria;p__Abditibacteriota;c__Abditibacteria;o__Abditibacteriales
d__Bacteria;p__Proteobacteria;c__Gammaproteobacteria;o__Piscirickettsiales
d__Bacteria;p__Bacteroidota;c__Bacteroidia;o__ML602M-17
d__Bacteria;p__Proteobacteria;c__Alphaproteobacteria;o__Alphaproteobacteria_Incertae_Sedis
d__Bacteria;p__Firmicutes;c__Bacilli;o__Thermoactinomycetales
d__Bacteria;p__Proteobacteria;c__Alphaproteobacteria;o__Caedibacterales
d__Bacteria;p__Nitrospirota;c__Nitrospiria;o__Nitrospirales
d__Bacteria;p__Firmicutes;c__Moorellia;o__Moorellales
d__Bacteria;p__Bacteroidota;c__Chlorobia;o__Chlorobiales
d__Bacteria;p__Desulfobacterota;__;__
d__Bacteria;p__Gemmatimonadota;c__Longimicrobia;o__Longimicrobiales
d__Bacteria;p__Firmicutes;c__Clostridia;o__Peptococcales
d__Bacteria;p__Proteobacteria;c__Gammaproteobacteria;o__Aeromonadales
d__Bacteria;p__Firmicutes;c__Clostridia;o__Clostridia
d__Bacteria;p__Actinobacteriota;c__Actinobacteria;o__Euzebyales
d__Bacteria;p__Firmicutes;c__Negativicutes;o__Veillonellales-Selenomonadales
d__Bacteria;p__Acidobacteriota;c__Acidobacteriae;o__Acidobacteriae
d__Bacteria;p__Firmicutes;c__Bacilli;o__RF39
d__Bacteria;p__Firmicutes;c__Thermaerobacteria;o__Thermaerobacterales
d__Bacteria;p__Verrucomicrobiota;c__Verrucomicrobiae;o__Verrucomicrobiales
d__Bacteria;p__Desulfobacterota;c__Desulfuromonadia;o__Desulfuromonadia
d__Bacteria;p__Planctomycetota;c__vadinHA49;o__vadinHA49
d__Bacteria;p__Chloroflexi;c__Chloroflexia;o__Thermomicrobiales
d__Bacteria;p__Planctomycetota;c__Phycisphaerae;o__C86
d__Bacteria;p__Verrucomicrobiota;c__Verrucomicrobiae;__
d__Bacteria;p__Proteobacteria;c__Gammaproteobacteria;o__Chromatiales
d__Bacteria;p__Patescibacteria;c__Gracilibacteria;__
d__Bacteria;p__Acidobacteriota;c__Blastocatellia;o__Blastocatellales
d__Bacteria;p__Actinobacteriota;c__Actinobacteria;o__Streptosporangiales
d__Bacteria;p__Dadabacteria;c__Dadabacteriia;o__Dadabacteriales

d__Bacteria;p__Planctomycetota;c__Phycisphaerae;__
d__Bacteria;p__Proteobacteria;c__Gammaproteobacteria;o__B2M28
d__Bacteria;p__Actinobacteriota;__;__
d__Bacteria;p__Desulfobacterota;c__Desulfobacteria;o__Desulfobacterales
d__Bacteria;p__Patescibacteria;c__Gracilibacteria;o__Candidatus_Peregrinibacteria
d__Bacteria;p__Proteobacteria;c__Gammaproteobacteria;o__Milano-WF1B-44
d__Bacteria;p__Actinobacteriota;c__Rubrobacteria;o__Rubrobacterales
d__Bacteria;p__Entotheonellaeota;c__Entotheonellia;o__Entotheonellales
d__Bacteria;p__Verrucomicrobiota;c__Omnitrophia;o__Omnitrophales
d__Bacteria;p__Proteobacteria;c__Gammaproteobacteria;o__F9P41300-M23
d__Bacteria;p__Acidobacteriota;c__Acidobacteriae;o__PAUC26f
d__Bacteria;p__Verrucomicrobiota;c__Lentisphaeria;o__P.palmC41
d__Bacteria;p__Proteobacteria;c__Magnetococcia;o__Magnetococcales
d__Bacteria;p__Proteobacteria;c__Alphaproteobacteria;o__AT-s3-44
d__Bacteria;p__Acidobacteriota;c__Holophagae;o__Holophagales
d__Bacteria;p__Planctomycetota;__;__
d__Bacteria;p__Planctomycetota;c__Planctomycetes;o__Gemmatales
d__Bacteria;p__Acidobacteriota;c__Subgroup_5;o__Subgroup_5
d__Bacteria;p__Bacteroidota;c__Ignavibacteria;o__Ignavibacteriales
d__Bacteria;p__Patescibacteria;c__Parcubacteria;o__Candidatus_Adlerbacteria
d__Bacteria;p__Firmicutes;c__Negativicutes;__
d__Bacteria;p__Firmicutes;c__Negativicutes;o__Acidaminococcales
d__Bacteria;p__Proteobacteria;c__Gammaproteobacteria;o__BD7-8
d__Bacteria;p__Deinococcota;c__Deinococci;o__Deinococcales
d__Bacteria;p__Elusimicrobiota;c__Elusimicrobia;o__Elusimicrobiales
d__Bacteria;p__Synergistota;c__Synergistia;o__Synergistales
d__Bacteria;p__Chloroflexi;c__Ktedonobacteria;o__C0119
d__Bacteria;p__Planctomycetota;c__Phycisphaerae;o__mle1-8
d__Bacteria;p__Cyanobacteria;c__Sericytochromatia;o__Sericytochromatia
d__Bacteria;p__Verrucomicrobiota;c__Verrucomicrobiae;o__Arctic97B-4_marine_group
d__Bacteria;p__RCP2-54;c__RCP2-54;o__RCP2-54
d__Bacteria;p__LCP-89;c__LCP-89;o__LCP-89
d__Bacteria;p__Elusimicrobiota;c__Lineage_IIb;o__Lineage_IIb
d__Bacteria;p__Bacteroidota;__;__
d__Bacteria;p__WPS-2;c__WPS-2;o__WPS-2

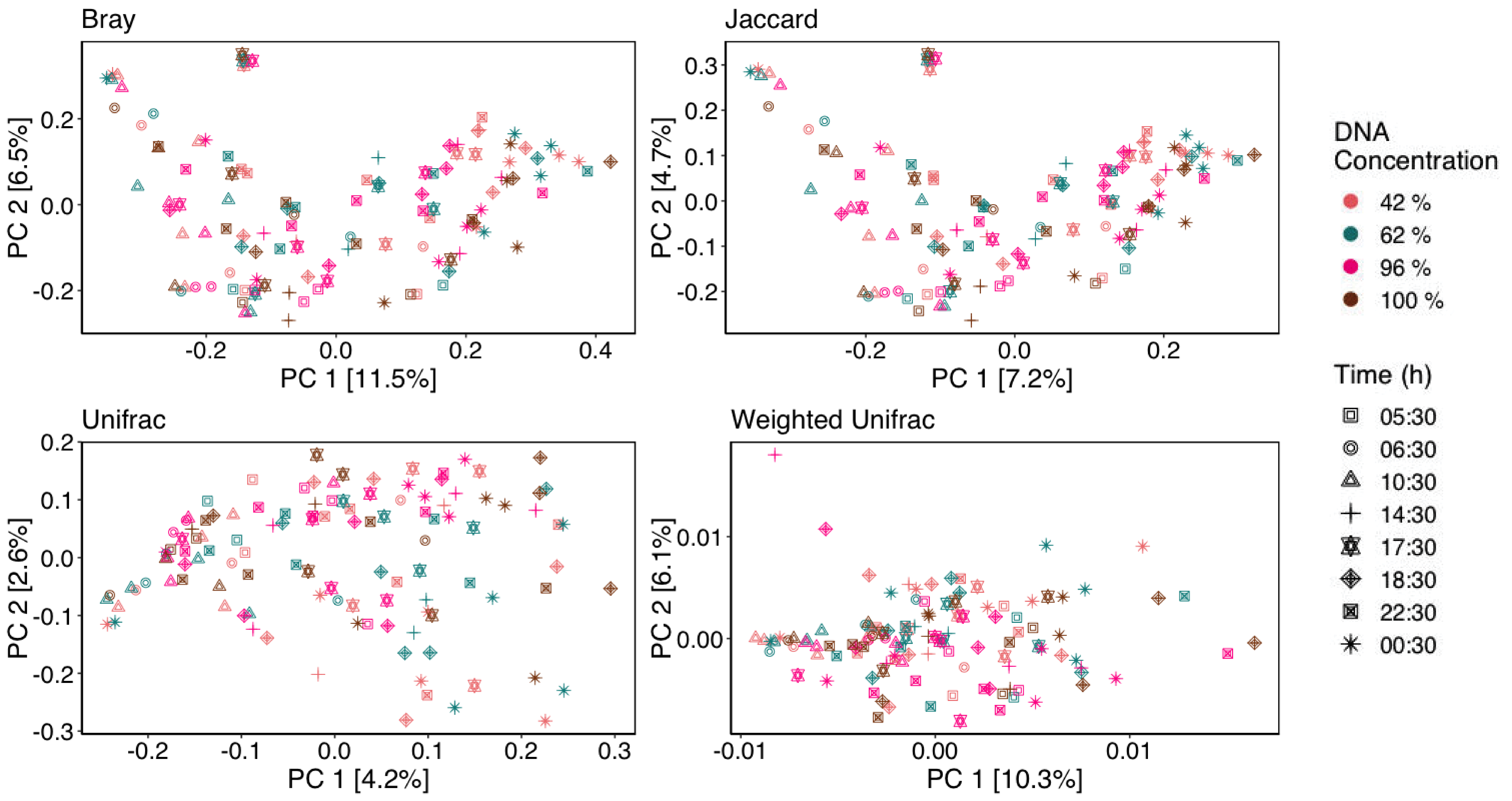
Bray
PC 2 [6.5%]
PC 1 [11.5%]
Jaccard
PC 2 [4.7%]
PC 1 [7.2%]
Unifrac
PC 2 [2.6%]
PC 1 [4.2%]
Weighted Unifrac
PC 2 [6.1%]
PC 1 [10.3%]
DNA Concentration
42 %
62 %
96 %
100 %
Time (h)
05:30
06:30
10:30
14:30
17:30
18:30
22:30
00:30
